# Benchmarking and Experimental Validation of Machine Learning Strategies for Enzyme Engineering

**DOI:** 10.64898/2026.03.29.715152

**Authors:** Zishuo Zeng, Jiao Jin, Rufang Xu, Xiaozhou Luo

## Abstract

Enzyme-directed evolution increasingly relies on computational tools to prioritize mutations, yet their practical value is difficult to assess because kinetic data are often aggregated across heterogeneous assay conditions, inflating apparent generalization. Here we introduce EnzyArena, a curated benchmark in which kinetic parameters (kcat, Km and kcat/Km) are grouped by experimental campaign and measured under matched conditions, thereby minimizing confounders such as pH, temperature, reaction, and batch. Using EnzyArena, we systematically evaluate 20 models spanning four widely used strategies for guiding enzyme evolution: protein–ligand binding affinity prediction, protein stability prediction, zero-shot fitness prediction and enzyme kinetic parameter prediction. Across subsets derived from public databases and 25 independent mutagenesis datasets, most models exhibit weak, fragile or inconsistent correlation with catalytic activity. Kinetic-parameter predictors perform strongly on database-derived subsets but lose their advantage on independent datasets, whereas zero-shot predictors show more consistent generalization. A simple consensus of multiple zero-shot models further improves the precision of identifying beneficial mutants. We prospectively validated these findings in a wet-lab campaign (150 mutants) comparing random mutants, UniKP-prioritized mutants and ESM-1v-prioritized mutants (representing supervised kinetic-parameter prediction and zero-shot fitness prediction, respectively), where ESM-1v achieved the highest utility and UniKP underperformed the random baseline. Together, this study establishes realistic baselines for computational mutant prioritization and highlights consensus zero-shot strategies as a practical starting point for enzyme engineering.

Zero-shot fitness predictors are the most competitive and generalizable class, but still achieve only modest correlations on independent datasets. Notably, a simple consensus of multiple zero-shot models substantially improves the precision of identifying superior mutants and currently offers the most practical computational benefit for enzyme engineering. Our work establishes EnzyArena as a fair benchmark for enzyme activity prediction, defines realistic performance baselines for existing tools, and highlights consensus zero-shot strategies as the most useful computational tool in practice for enzyme-directed evolution.

## Introduction

Enzyme-directed evolution underpins modern biocatalysis, green chemistry ^1^and industrial biotechnology ^2^ by enabling the rapid optimization of catalysts for non-native substrates, process conditions and reaction networks. Over the past few years, advances in machine learning have transformed how many groups design and prioritize variants ^3^, with tools ranging from protein language models ^4–7^ and neural-network kinetic predictors ^8–12^ to docking-based binding models ^13–17^ and stability scoring predictors ^18–22^ now routinely appearing in high-profile enzyme engineering studies. In principle, these approaches promise to compress the search space, reduce the experimental burden ^23^ and reveal general principles of sequence–function relationships that are difficult to infer from intuition alone.

Yet the current empirical foundation for these promises is surprisingly fragile. Many published studies ^8–12^ aggregate kinetic data obtained under heterogeneous assay conditions, differing in temperature, pH, substrate identity, cofactors, host background and assay format, and then treat the resulting *kcat*, *Km* or *kcat*/*Km* values as if they were directly comparable. Endpoints with distinct biophysical meanings, such as catalytic turnover, ligand binding, protein stability and organismal growth, are often analyzed interchangeably, with activity, “fitness” and expression level being used as proxies for one another. On top of this, enzymology itself is marked by substantial metrological variation: even when nominal assay conditions and protocols are matched, measurements from different laboratories, different instruments or even two nominally identical devices can diverge ^24^ because of differences in calibration, baseline correction, and timing. Machine-learning models trained on such aggregates are therefore encouraged to fit a mixture of sequence–function signal and instrument- or laboratory-specific biases. Finally, models trained on large public resources such as BRENDA ^25^ and SABIO-RK ^26^ are frequently evaluated on test sets that partially overlap with their training data, or that share closely related sequences and experimental campaigns, leading to hidden data leakage and optimistic performance estimates ^27^ ^28^ ^29^. Together, assay heterogeneity, endpoint inconsistency, metrological variability and data leakage make it difficult to know which computational strategies genuinely enrich for variants with improved catalytic performance and which succeed mainly by exploiting quirks of current datasets.

In this work, we address this problem from a data-centric perspective by constructing a curated benchmark (namely, EnzyArena) for computationally guided enzyme-directed evolution built around kinetic parameters measured under strictly matched conditions and grouped at the level of individual experimental campaigns. We assemble datasets in which temperature, pH, substrate, host and assay format are controlled within each subset, explicitly track potential overlaps with public kinetic databases, and treat laboratory- and instrument-specific effects as fundamental sources of variation rather than negligible noise (Figure 1A). On this foundation, we systematically compare four major families of models—binding predictors, stability scorers, zero-shot protein language models and kinetic-parameter predictors (Figure 1B)—using evaluation protocols designed to avoid leakage and to mirror realistic directed-evolution use cases. Under these stringent conditions, we find that many widely used models show limited and fragile correspondence with catalytic outcomes, whereas zero-shot sequence models provide the most robust and transferable signal for ranking enzyme variants. We then complement this retrospective benchmarking with a prospective wet-lab enzyme engineering campaign to directly test whether model-guided recommendations improve mutant prioritization under practical experimental constraints.

**Figure 1.**
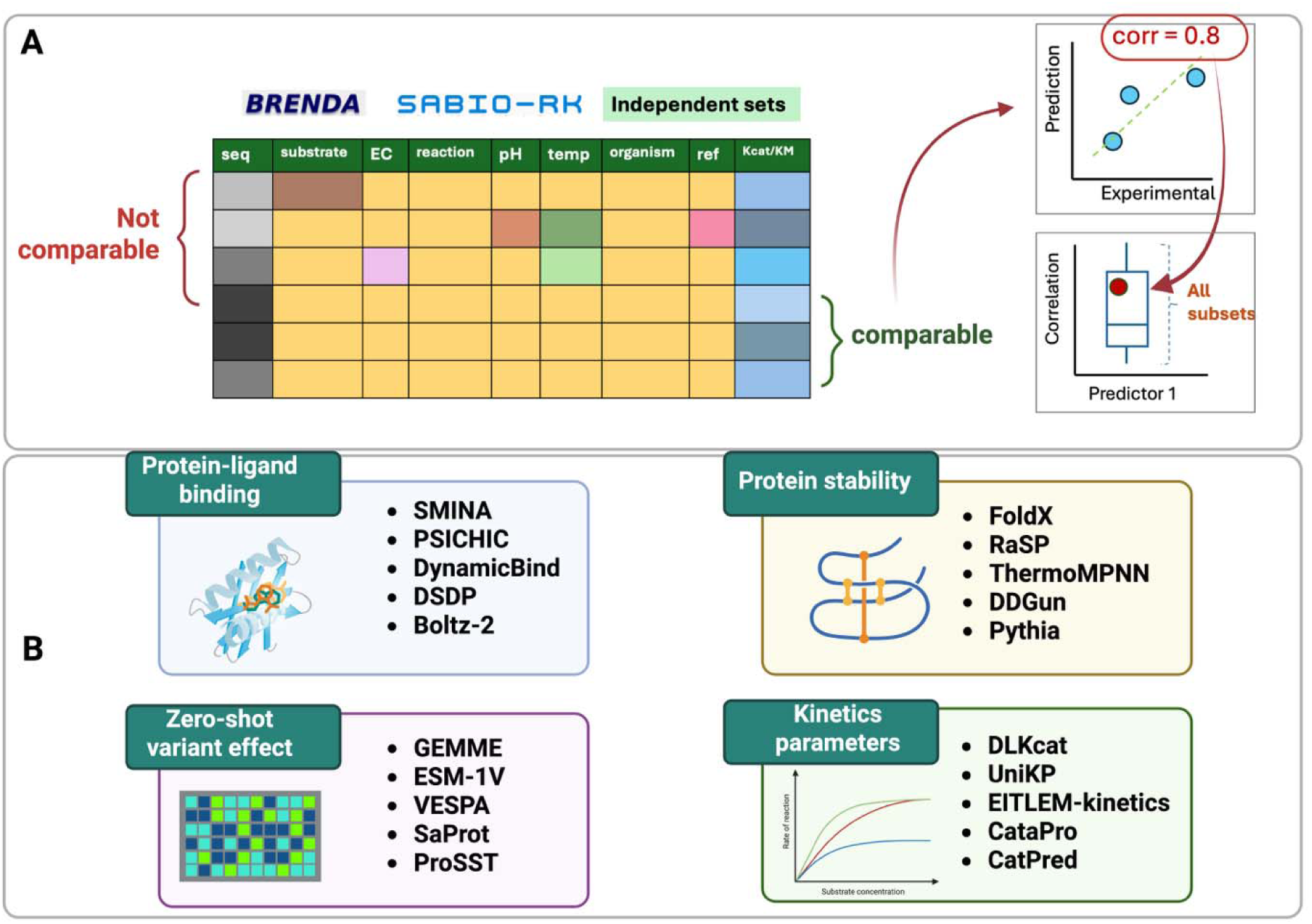
Overview of the methodology in this study. Panel A shows the data cleansing process and method of evaluation. Data of enzyme KP were obtained from public databases and organized into multiple subsets where all experimental conditions are consistent across samples within subset. Predictions from various computational tools were made for each of the subsets and the Spearman correlations for all the subsets were computed and summarized for each individual predictor. Panel B summarized the computational tools included in this study for four different commonly used computational strategies, including protein-ligand binding affinity prediction, protein stability prediction, zero-shot fitness prediction, and enzyme KP prediction.

## Results and Discussion

### Data curation

A fair evaluation of computational enzyme predictors requires a dataset that minimizes experimental confounders while maintaining broad coverage across enzyme classes. We argue that an ideal benchmarking dataset should meet two key criteria: (1) it should encompass a wide range of enzyme types to reduce functional bias, and (2) experimental conditions—such as pH, temperature, and expression system—should remain consistent within subsets to avoid confounding factors. However, many existing kinetic datasets lack detailed metadata describing assay conditions, which limits reproducibility and cross-study comparability ^30^. In particular, the training of current kinetic-parameter (KP) predictors often violates the second criterion, as they aggregate data from heterogeneous sources during training. Likewise, while repositories such as ProteinGym provide large-scale fitness datasets with diverse experimental labels, including binding, organismal fitness, expression, stability, and activity, the majority of activity data involve non-enzymatic functions or enzyme reactions with macromolecular substrates, which contradicts with KP predictors in this study, as they support only small-molecule substrates. These limitations motivated our effort to construct condition-controlled datasets suitable for rigorous benchmarking.

Following these principles, we collected 1,571 *kcat* (turnover number), 2,250 *Km* (Michaelis constant), and 1,351 *kcat*/*Km* (catalytic efficiency) entries from the BRENDA and SABIO-RK databases, applying strict inclusion criteria (i.e., mutagenesis studies with ≥10 variants per assay, including one wild-type and at least nine mutants; see Materials and Methods). The curated data were partitioned into 130 *kcat* subsets, 174 *Km* subsets, and 110 *kcat*/*Km* subsets, ensuring internal consistency across experimental parameters such as pH, temperature, source organism, substrate, reaction type, and reference. This structured partitioning ensured that data within each subset were comparable and that observed performance differences among predictors could be attributed to algorithmic rather than experimental variation.

In addition to the KP datasets, we manually curated 25 independent datasets related to enzyme catalytic activity that are not included in BRENDA or SABIO-RK. These comprise four deep mutagenesis scanning (DMS) fitness score datasets, nine *kcat* datasets, nine *Km* datasets, and three *kcat*/*Km* datasets (Table 1). These independent datasets were particularly important for unbiased evaluation of kinetic-parameter predictors, which are frequently trained on data derived from BRENDA and SABIO-RK. Together, these curated and independent datasets form the foundation of **EnzyArena**, enabling fair and reproducible assessment of enzyme activity prediction models under well-controlled experimental conditions.

**Table 1.**
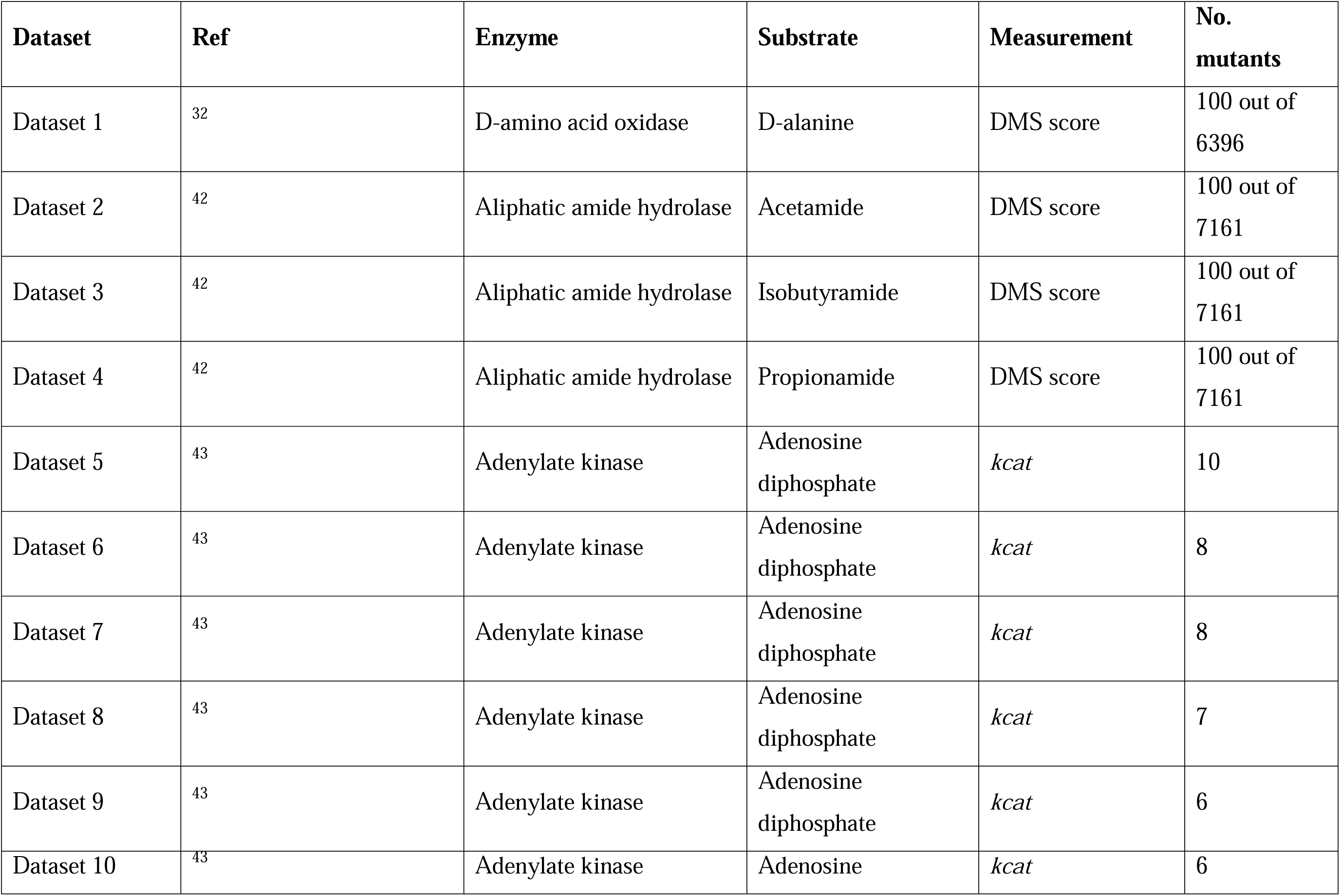
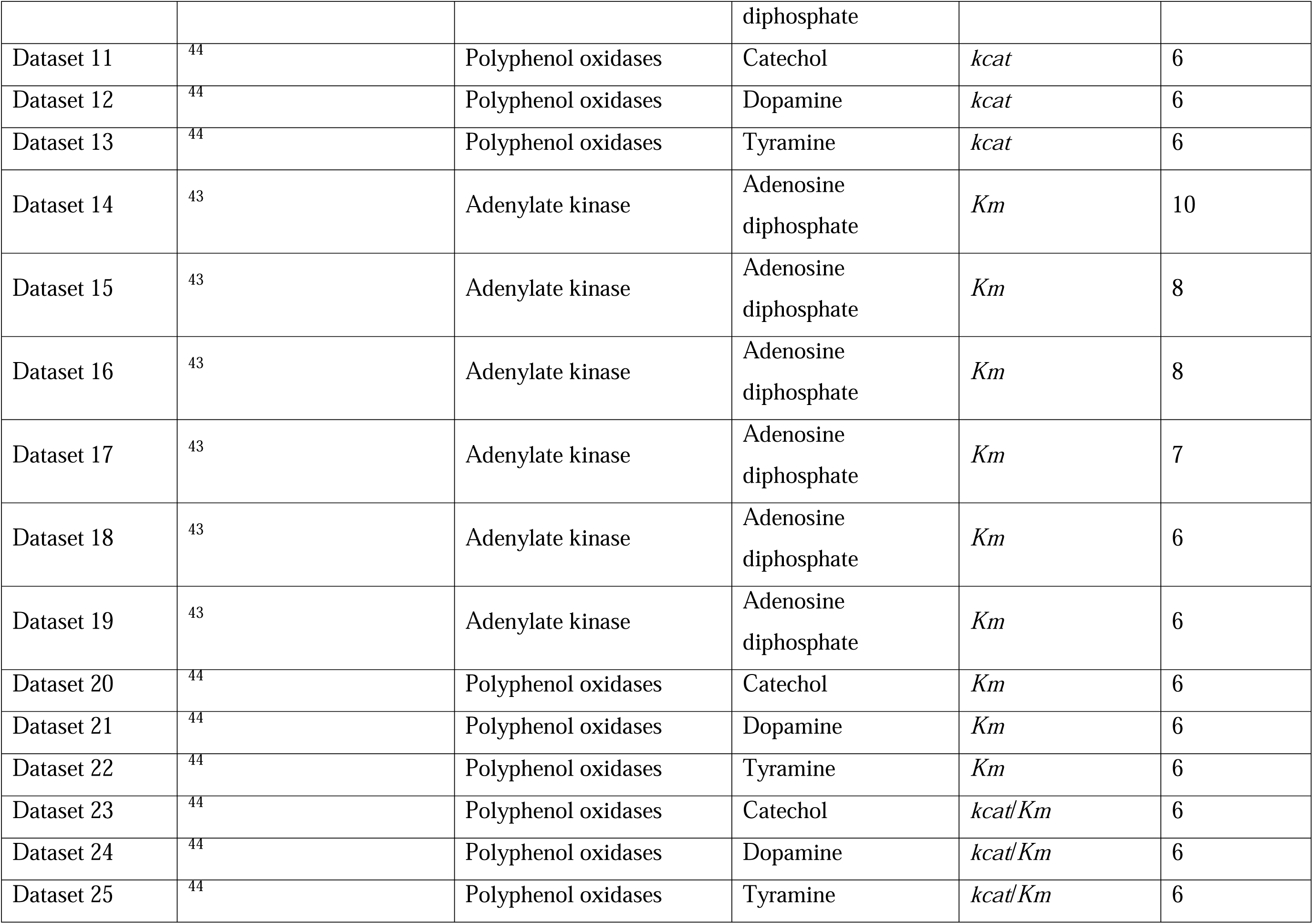
Summary of the manually curated datasets.

### Selection of Representative Predictors for Each Computational Strategy

To provide a fair and comprehensive evaluation of commonly used computational strategies for enzyme engineering, we selected five representative tools for each of the four major paradigms examined in this study: protein–ligand binding affinity prediction, protein stability prediction, zero-shot fitness prediction, and kinetic-parameter (KP) prediction (Table 2). Selection of predictors are based on the considerations of methodological diversity, widespread use, recency of publication, and performance in independent benchmark when available. Detailed descriptions of the individual models can be found in Supplementary Information.

**Table 2.**
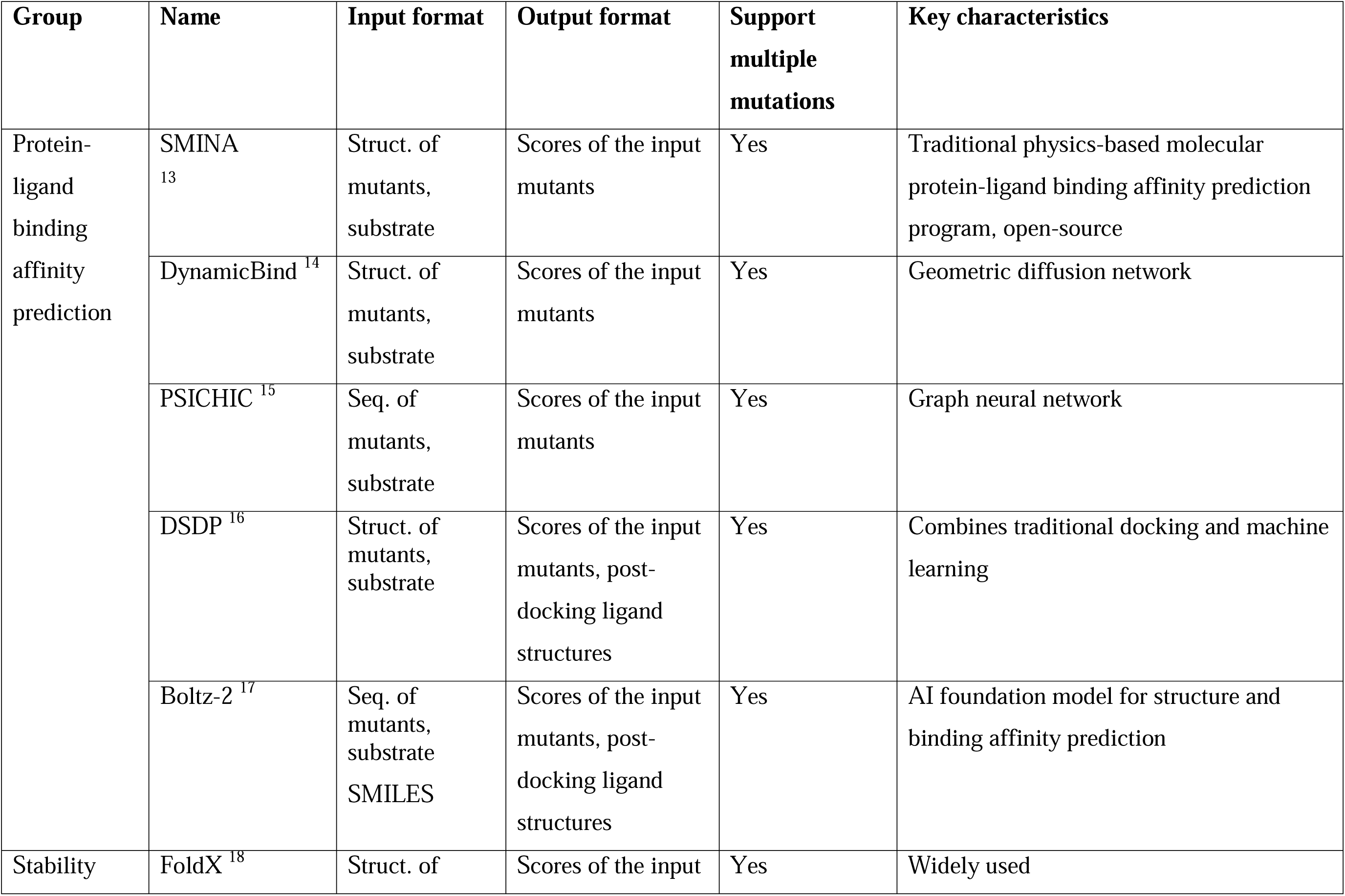
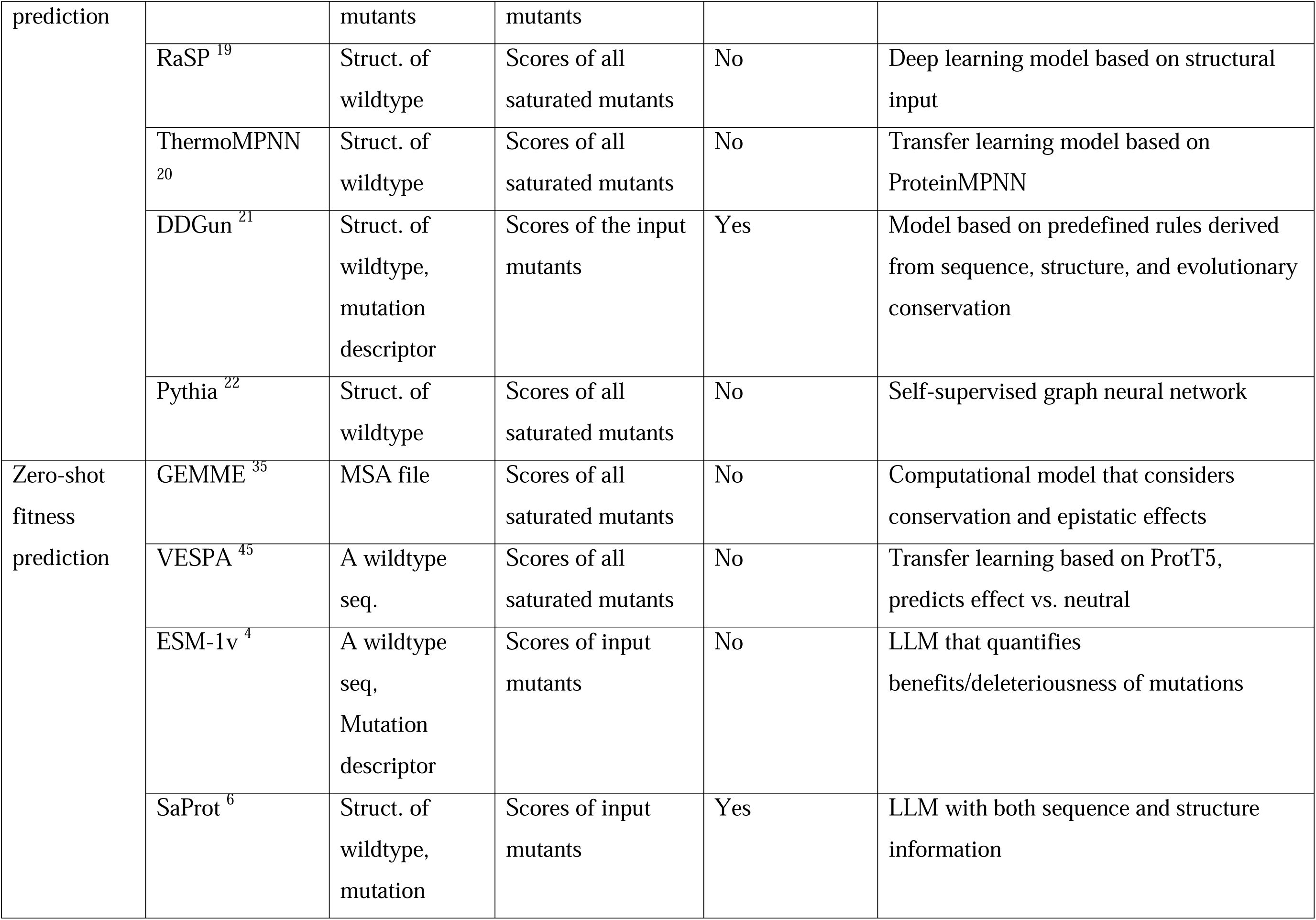
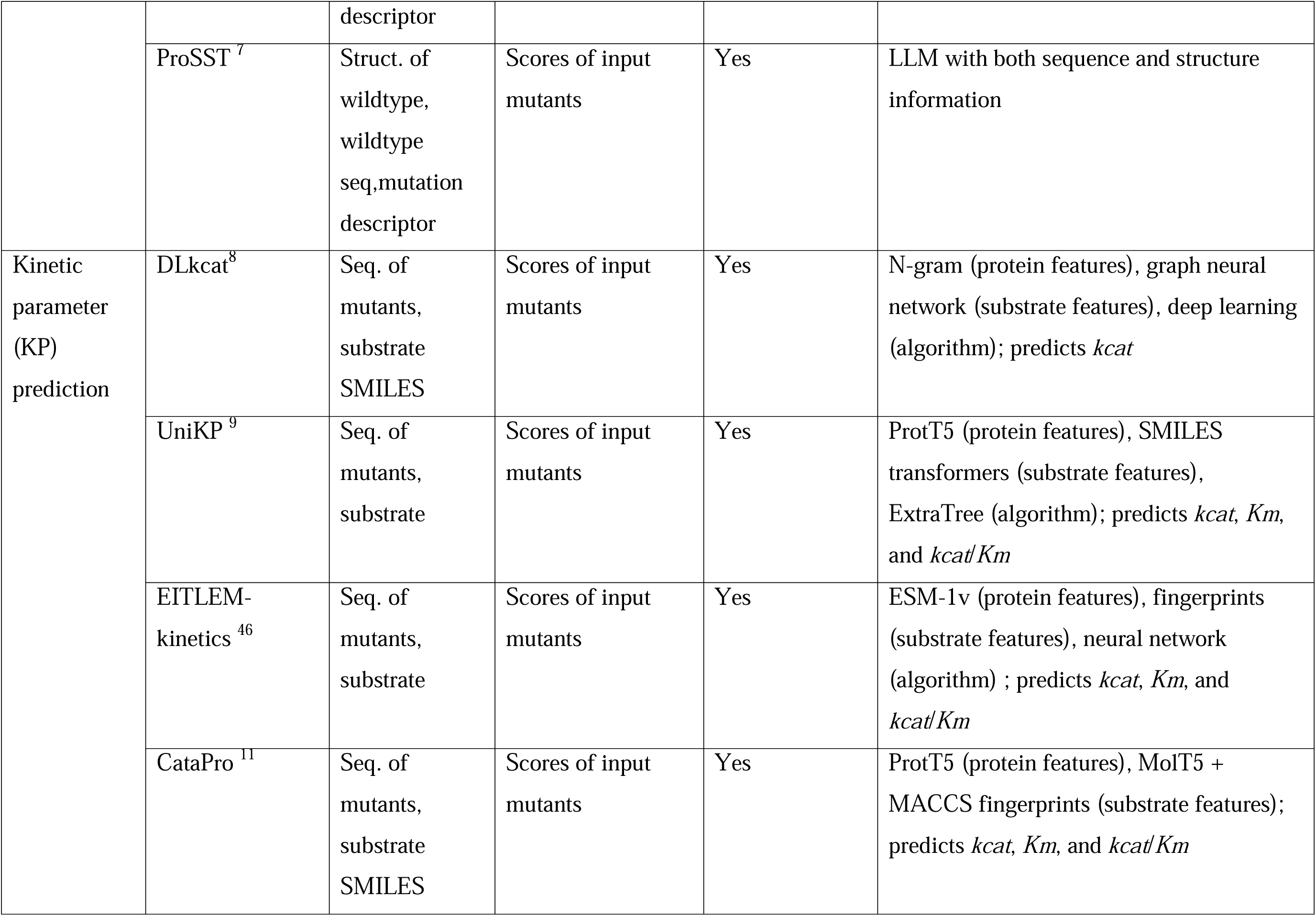
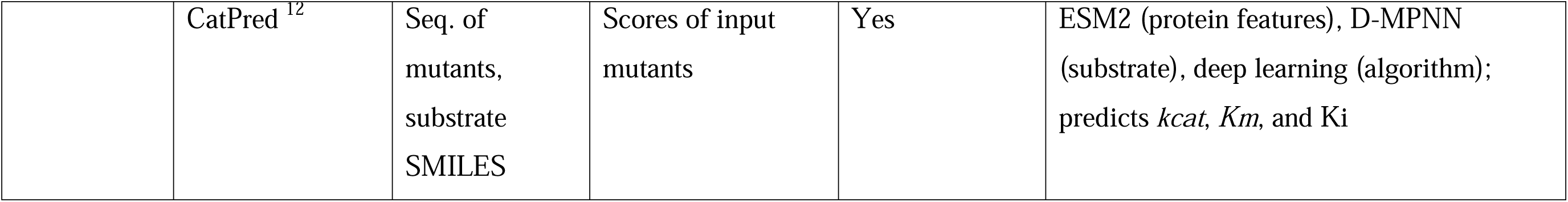
Summary of the selected predictors in this study.

Although stability, binding affinity, and zero-shot fitness are not mechanistically equivalent to catalytic activity, they are nevertheless widely used proxies in enzyme-directed evolution for several practical reasons. First, protein stability is often treated as a prerequisite for high activity and evolvability: stabilizing mutations can expand the sequence space accessible to laboratory evolution and mitigate activity–stability trade-offs ^31–33^. Second, binding-affinity predictions, particularly docking scores, are frequently used to identify mutations that strengthen substrate interactions, with numerous studies reporting improved catalytic performance after selecting variants with improved predicted binding ^34^. Third, zero-shot fitness predictors have gained traction due to their accessibility and scalability due to their demonstrated utility in identifying beneficial or deleterious mutations across large mutational landscapes, including in enzymes ^30^ ^4^ ^35, 36^. This widespread use underscores the need for a systematic, leakage-free, and condition-controlled evaluation, thus motivating the benchmarking framework established in this study.

Table 3 reports the computational time required to run all models on a compute node of NVIDIA A10 GPU with 32 CPUs. Most stability predictors and kinetic parameter (KP) predictors completed within a short timeframe, with the exception of FoldX and EITLEM-Kinetics, which required 17.83 hours and 8 hours, respectively. All zero-shot predictors were computationally efficient, completing the full dataset in under 5 hours. In contrast, binding affinity prediction tools were substantially more time-consuming. Except for PSICHIC, all docking-based methods required between 31.5 and 112 hours to process the entire dataset, making them the slowest among the evaluated model categories.

**Table 3.** Model time consumption to run all data in this study.

| Category | Model | Data processed | Model run time (h) | Run time for pdb/pdbqt preparation (h) | Total run time (h) |
| --- | --- | --- | --- | --- | --- |
| Stability | Pythia | 126 wildtype protein pdb files | 0.05 | 1 | 1.05 |
|  | ThermoMPN N |  | 0.17 | 1 | 1.17 |
|  | DDGun |  | 1.17 | 1 | 2.17 |
|  | RaSP |  | 5 | 1 | 6 |
|  | FoldX | 2195 protein pdb files | 0.83 | 17 | 17.83 |
| Kinetic Parameters | DLKcat | 2886 protein sequence - substrate pairs | 0.08 | - | 0.08 |
|  | UniKP |  | 0.5 | - | 0.5 |
|  | CataPro |  | 0.5 | - | 0.5 |
|  | CatPred |  | 0.5 | - | 0.5 |
|  | EITLEM-kinetics |  | 8 | - | 8 |
| Fitness | SaProt | 126 wildtype protein pdb files | 0.33 | 1 | 1.33 |
|  | ESM1V |  | 2 | - | 2 |
|  | ProSST |  | 1 | 1 | 2 |
|  | GEMME |  | 2.5 | - | 2.5 |
|  | VESPA |  | 4.33 | - | 4.33 |
| Binding Affinity | PSICHIC | 2886 protein sequence - substrate pairs | 0.17 | - | 0.17 |
|  | DSDP |  | 1 | 30.5 | 31.5 |
|  | SMINA |  | 16 | 30.5 | 46.5 |
|  | DynamicBin<br>d |  | 85 | 17 | 102 |
|  | Boltz-2 |  | 112 | - | 112 |

### Zero-shot Fitness Predictors Outperformed Binding Affinity and Stability Predictors for All Data Sources

To begin benchmarking the different computational strategies, we first compared stability, binding-affinity, and zero-shot fitness predictors across all three data sources, including BRENDA, SABIO-RK, and the manually curated independent datasets—because none of these model families were trained on these datasets. This ensures that all comparisons in this section are free from data leakage and reflect each model’s intrinsic generalizability. An overview of model performance on BRENDA- and SABIO-RK–derived subsets is shown in Figure 2 (with detailed breakdowns in Figures S1 and S2).

**Figure 2.**
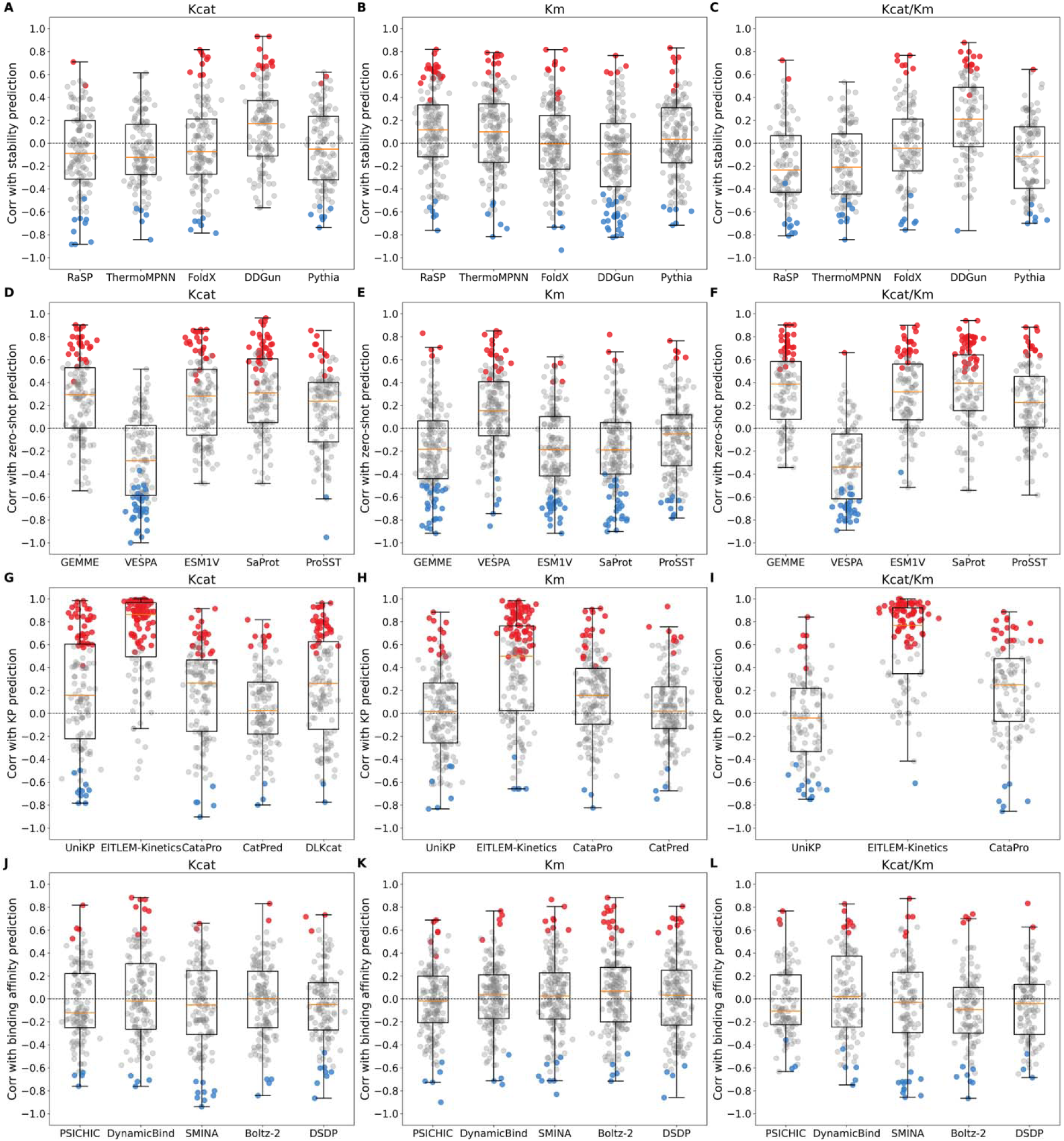
Performances overview of computational tools on subsets derived from BRENDA and SABIO-RK. Performance is measured by Spearman correlation. In each panel, red, blue, and grey dots indicate significantly (p-value < 0.05) positive, significantly negative, and non-significant Spearman correlations, respectively.

Across all data sources, zero-shot fitness predictors consistently achieved the highest correlations with experimental measurements. Their median Spearman correlations ranged from 0.134 to 0.254 in BRENDA subsets (Figure S1), 0.173 to 0.377 in SABIO-RK subsets (Figure S2), and 0.15 to 0.25 overall—BRENDA, SABIO-RK and independent datasets (Figure 3A). This trend also holds when sample size is accounted (Figure 3B).

**Figure 3.**
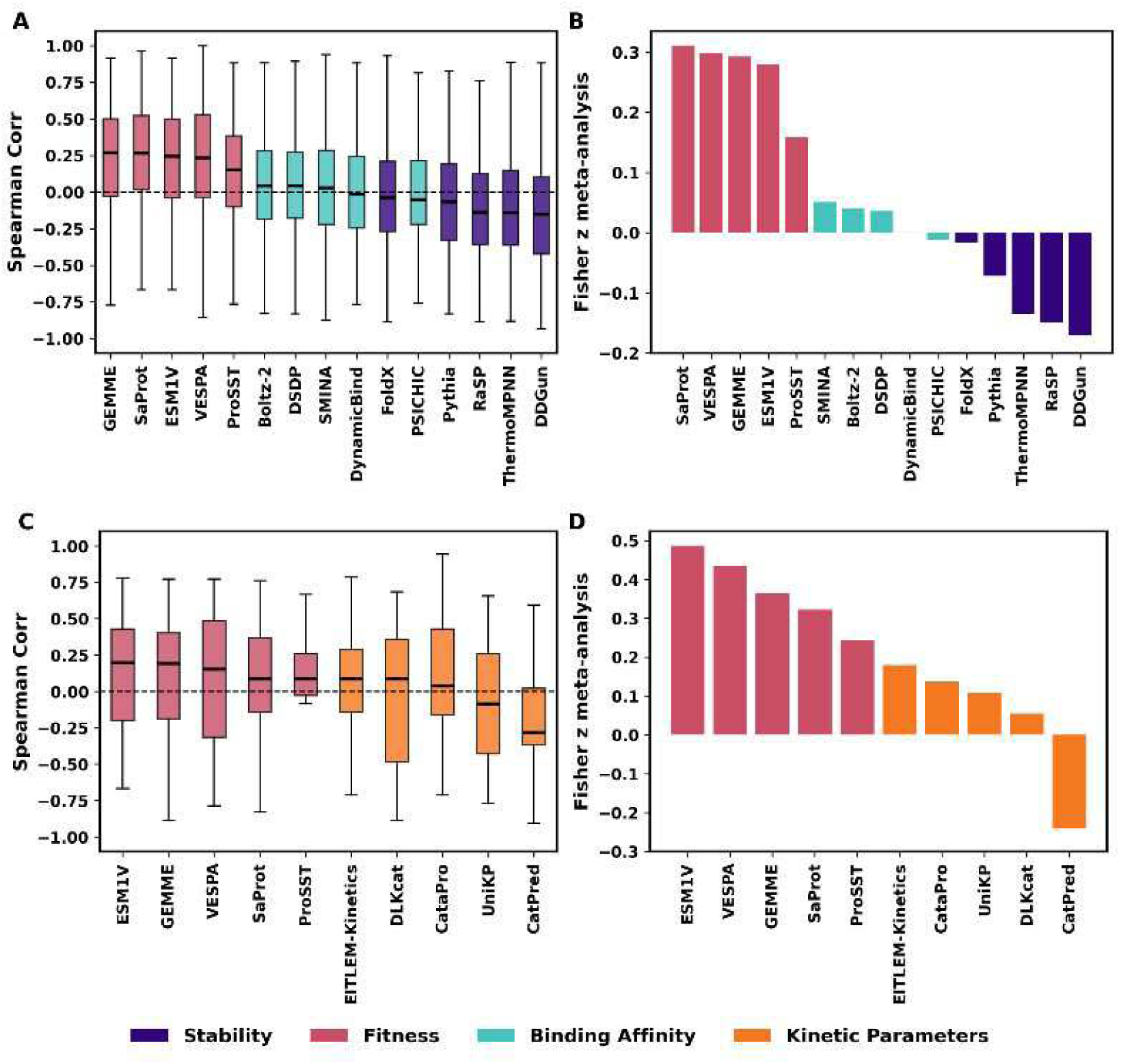
Comparative performance evaluation of computational predictors across datasets. (A–B) Comparison among stability, zero-shot fitness, and binding-affinity predictors across all three data sources (BRENDA, SABIO-RK, and independent datasets). (C–D) Comparison between zero-shot fitness and kinetic-parameter predictors using only independent datasets. Panels A and C show the distribution of Spearman correlation coefficients across subsets, ordered by median correlation from highest to lowest. Panels B and D summarize the weighted sum of correlations (weighted by sample size, Eqn. 3), also ordered from highest to lowest. To ensure consistent interpretation, correlation signs were adjusted by taking absolute values where appropriate to match the expected relationship with DMS scores, *kcat*, *Km*, and *kcat*/*Km* (see Materials and Methods and Table S1), so that higher correlations uniformly indicate better model performance.

In contrast, binding-affinity predictors showed near-zero association with experimental kinetic parameters. For example, correlations with Km were minimal, with median values between – 0.017 and 0.038 (Figure 2K). Although *Km* is often interpreted as a proxy for substrate binding affinity, in theory, *Km* reflects a composite of binding affinity (Kd), catalytic turnover (*kcat*), and the rates of substrate association and dissociation ^37^. Moreover, solvent effects such as pH and ionic strength, which critically modulate enzyme–substrate interactions, are typically not accounted for in computational binding models ^38^. Therefore, binding-affinity predictions alone are insufficient to capture the multifactorial determinants of experimental *Km*, which may partially explain the underperformances of binding affinity predictors on *Km* datasets.

Similarly, protein stability predictions showed little to no correlation with experimental measurements (with medians of correlations ranging from −0.018 to 0.051; Figure 2A-C). Although stability and catalytic activity are theoretically independent properties, stability predictors are widely incorporated into protein engineering workflows under the assumption that enhanced stability may indirectly support improved activity ^33^, nevertheless, a recent benchmark study reported that predictive accuracy for stability remains limited ^39^. Consistent with these findings, our results indicate that current stability predictors provide limited utility in guiding catalytic activity optimization.

Together, these results show that evolutionary or language model–derived signals provide the most informative basis for ranking enzyme variants across diverse conditions, even though zero-shot models were never trained on catalytic activity labels.

### Kinetic Parameter Predictors Succeed on KP Datasets but Lose Advantage on Leakage-Free Independent Data

Because zero-shot predictors outperformed both stability and binding-affinity models across all datasets (Figure 3A), we next compared zero-shot predictors directly with kinetic-parameter (KP) predictors. KP models represent the most mechanistically relevant strategy for catalytic prediction, but they are typically trained on large volumes of BRENDA and SABIO-RK data. As a result, their performance on subsets derived from these databases is susceptible to data leakage. Indeed, in the BRENDA and SABIO-RK subsets, one of the KP predictors, EITLEM-kinetics, achieved the highest median Spearman correlations (median = 0.680), while the remaining KP models performed similarly to zero-shot predictors with modest correlations (0.024-0.263) (Figure 2). These strong performances are expected given the overlap in data sources and highlight the need for evaluations on leakage-free datasets.

To obtain an unbiased comparison, we therefore benchmarked both strategy classes on the manually curated independent datasets (Figure 3C), which share no overlap with the training data of KP predictors. On these truly independent datasets, both zero-shot and KP predictors exhibited reduced performance (Figure 3C), with zero-shot predictors reaching median correlations of 0.086-0.200 and KP predictors reaching −0.283-0.086. These modest correlations reflect two sources of limitation: a) model limitations, where predictors fail to generalize outside the domain of their training data; and b) data limitations, as many of the independent datasets contain only 6–10 variants. Such small sample sizes amplify technical noise and bias, making rank-based metrics like Spearman correlation particularly unstable ^40^.

For a more robust summary that accounts for differences in dataset size, we combined correlation estimates using Fisher z meta-analysis ^41^. In this framework, correlations are transformed to stabilize variance and then pooled using inverse-variance weighting so that larger and more informative datasets contribute more strongly to the overall estimate (Eqn. 3; Materials and Methods). This approach is widely used to aggregate correlation coefficients across heterogeneous studies and is particularly appropriate when sample sizes differ substantially between datasets {Borenstein, 2021 #1318}. When correlations were weighted by dataset size (Figure 3D), zero-shot predictors still retained an advantage (weighted sum = 0.219 to 0.406) over KP models (weighted sum = −0.197 to 0.157), reflecting slightly stronger generalization despite the noisy, small-scale evaluation setting.

Overall, these results show that while KP predictors showed good perform on KP datasets overlapped with their training sources, their advantage rapidly diminishes when evaluated on independent, leakage-free data. In contrast, zero-shot predictors maintain relatively robust performance across all conditions, suggesting that sequence-level evolutionary information confer stronger generalizability than current supervised KP prediction models.

### Inter-Model Correlation Analysis Reveals Strategy-Specific Clustering

To better understand relationships among predictors, we analyzed pairwise correlations across all model outputs (Figure 4). Models within the same computational paradigm formed clear clusters, with intra-group Pearson correlations ranging from −0.812 to 0.941, while inter-group correlations were generally weak (−0.518 to 0.485). Notable exceptions included: FoldX, which showed unusually low correlation with other stability predictors (correlation = −0.060 to 0.155), consistent with recent benchmarks noting its limited accuracy; KP predictors, which showed moderate correlation with binding-affinity predictors in the Km subset (correlation = −0.518 to 0.485), consistent with Km’s partial dependence on substrate binding. These structural relationships between models further support our strategy to evaluate predictors by family and to identify which paradigms generalize best across datasets.

**Figure 4.**
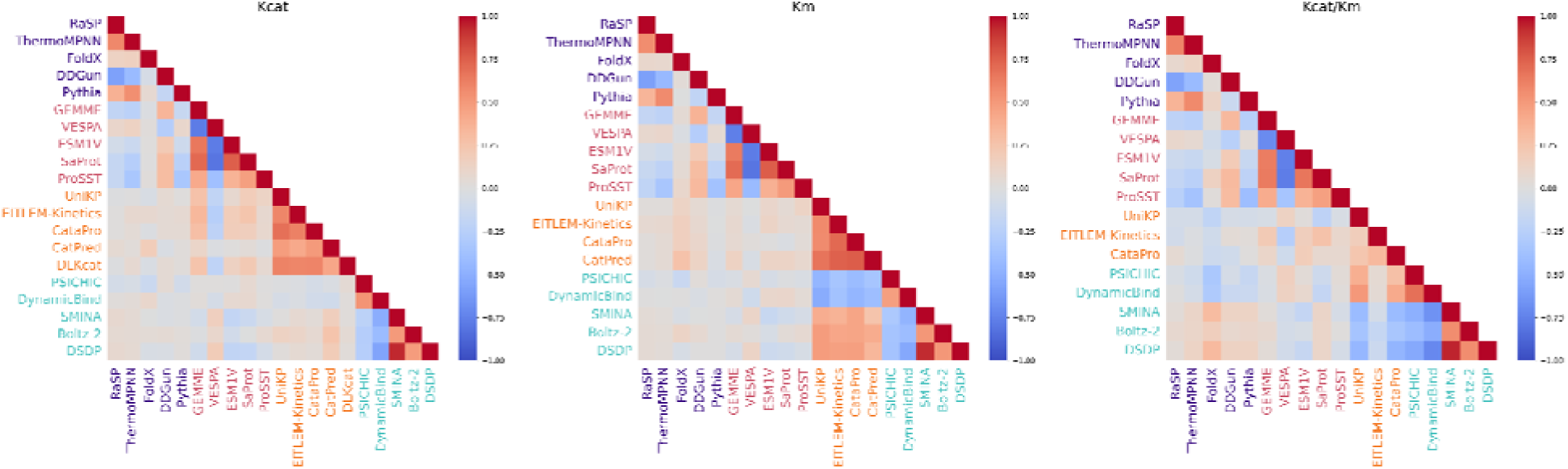
Heatmap of pairwise correlation among the predictors. The heatmaps display Pearson correlation coefficients between model predictions, with values ranging from +1 (red, perfect positive correlation) to –1 (blue, strong negative correlation). Models are grouped and color-coded based on computational strategy: purple (stability predictors), red (zero-shot sequence-based models), orange (kinetic parameter–specific models), and cyan (binding affinity models).

### Prospective wet-lab validation using patchoulol synthase

Building on the benchmarking results described above, two key conclusions emerge. First, zero-shot fitness predictors and kinetic-parameter (KP) predictors represent the strongest-performing strategies on current public datasets, with KP models achieving high apparent accuracy when evaluated on data sources closely related to their training distributions. Second, when evaluated on leakage-free independent datasets, zero-shot predictors consistently show better generalization than KP models, albeit under conditions where dataset sizes are typically small and correlations remain modest. Given that many of these independent datasets contain only a limited number of variants, it remains unclear whether the observed advantage of zero-shot models fully translates to practical enzyme engineering scenarios. To address this gap and directly assess real-world utility, we next performed a prospective wet-lab validation to compare model-guided mutation prioritization strategies under controlled experimental conditions.

To experimentally evaluate model-guided mutation prioritization, we selected patchoulol synthase (PTS) as a test enzyme. PTS catalyzes the cyclization of farnesyl pyrophosphate to patchoulol, which is the major bioactive component of patchouli oil and has attracted increasing attention due to its reported anticancer, neuroprotective, anti-inflammatory, and gastroprotective activities {Lee, 2020 #1319}. Despite its industrial relevance, PTS is a multiproduct terpene synthase that produces patchoulol at relatively low selectivity {Ekramzadeh, 2020 #1320}, limiting its biotechnological application and makes it a challenging target for enzyme engineering.

We performed saturated single-site mutagenesis on PTS and used computational models to prioritize variants for experimental testing. Based on the benchmarking results, ESM-1v and UniKP were selected to represent the zero-shot fitness strategy and KP strategy, respectively. Each model prioritized 50 mutants, and an additional 50 mutants were selected randomly as a baseline, resulting in 150 variants in total. All variants were constructed and experimentally evaluated by measuring patchoulol production relative to the wild type (Figure 5A).

**Figure 5.**
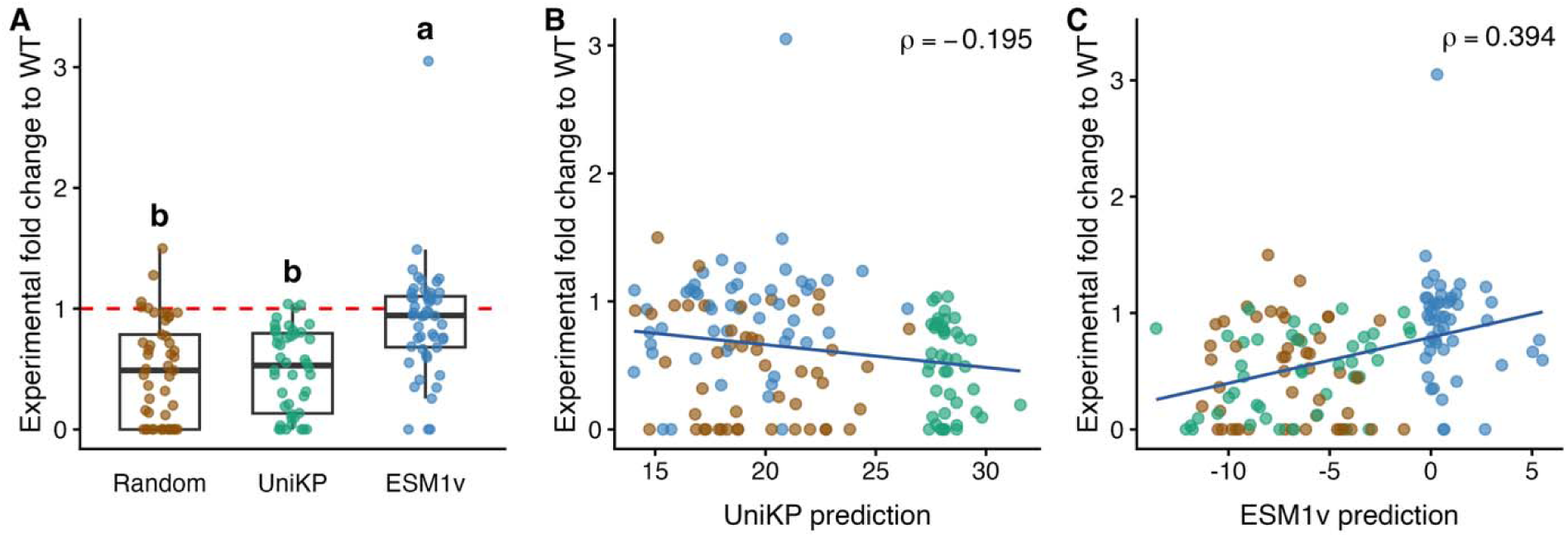
Model performance on a wet-lab validation dataset for model-guided mutagenesis of the PTS enzyme. (A) Distribution of experimentally measured fold change values for mutants prioritized by Random, UniKP, and ESM1v models. The red dashed line marks the wildtype (WT) level (fold change = 1). Letters above the boxes denote statistical groupings from a one-way ANOVA followed by Tukey’s post-hoc test; groups that do not share a letter differ significantly (P < 0.05). (B) Relationship between UniKP model predictions and experimentally measured fold change relative to wild type (WT). (B) Relationship between ESM1v model predictions and experimental fold change to WT. Each point represents a mutant, colored by model category; the Spearman correlation coefficient (ρ) is indicated. The regression line shows the least-squares fit. Together, these results show that ESM1v predictions exhibit a stronger monotonic relationship with experimental measurements compared with UniKP, and that model-guided designs yield distinct distributions of functional outcomes relative to random variants.

The experimental results show a clear separation between strategies. Variants prioritized by UniKP do not outperform the random baseline, whereas ESM-1v–prioritized variants show significantly higher activity than both random and UniKP-selected mutants (Figure 5A). To further examine model–experiment agreement, we pooled all 150 variants and performed a retrospective correlation analysis between model predictions and experimental measurements. UniKP predictions show a negative correlation with experimental activity (ρ = −0.195; Figure 5B), whereas ESM-1v predictions show a positive correlation (ρ = 0.394; Figure 5C). Notably, one ESM-1v–prioritized mutant (C23S) achieved an approximately threefold higher activity over the wild type.

Overall, this prospective validation supports the conclusions from the computational benchmark and demonstrates that zero-shot fitness prediction provides a more robust and generalizable strategy for prioritizing enzyme mutations in practice.

### Combination of Zero-Shot Predictors Improves the Precision of Identifying Superior Enzyme Mutants

Given the strong individual performance of zero-shot fitness predictors demonstrated above, we next investigated whether combining multiple predictors could further improve the success rate of identifying mutants with enhanced catalytic activity. Our analysis focused on precision (Eqn. 1), defined as the proportion of predicted beneficial mutants that are experimentally confirmed to outperform the reference. Because “wildtype” is context-dependent, we generated an exhaustive pairwise comparison dataset in which every mutant in turn served as the reference, and all other mutants were evaluated relative to it. This strategy greatly expanded the evaluation space and enabled a robust assessment of prediction performance across diverse mutational landscapes.

For each pairwise comparison, we evaluated whether a given predictor or combination of predictors correctly identified the mutant with superior experimental activity. To assess combinations, we implemented a consensus strategy, where a mutant was predicted to be superior only if all zero-shot models in the combination ranked the mutant higher than the reference. All 31 possible combinations of the five predictors (ranging from single models to combinations of all five) were evaluated (Figure 6A; Figure S1). Importantly, predictor outputs were directionally standardized so that higher scores consistently indicated higher predicted fitness.

**Figure 6.**
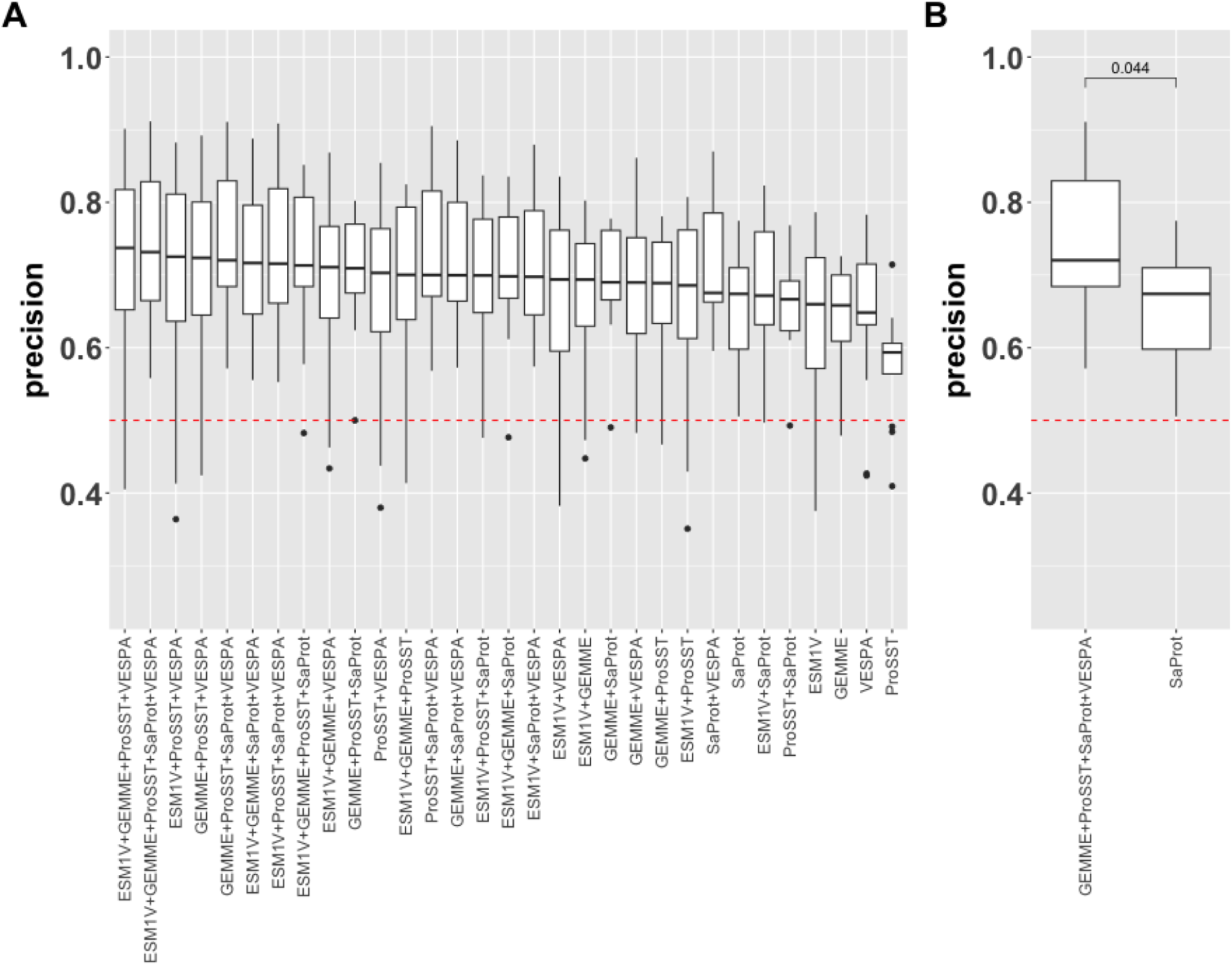
Combination of zero-shot predictors improves precision in identifying beneficial enzyme mutants. (A) Precision of all 31 combinations of five zero-shot predictors, evaluated using exhaustive pairwise comparisons across datasets with at least 20 mutants. Each box plot represents the distribution of precision values across datasets, with the dashed red line indicating random baseline performance. (B) Comparison of the best-performing single predictor (SaProt) and the top-performing combination (GEMME + ProSST + SaProt + VESPA). The consensus predictor achieves a significantly higher precision (mean 0.74) than SaProt alone (mean 0.65), demonstrating the benefit of integrating multiple predictive models (p = 0.044, Wilcoxon rank-sum test).

Across datasets containing at least 20 mutants, the combination of GEMME, ProSST, SaProt, and VESPA achieved a significantly higher average precision (0.74) compared to the best-performing single predictor, SaProt (0.65) (Wilcoxon rank-sum test, p = 0.044; Figure 6B). In contrast, for datasets with fewer than 20 mutants, all predictors, whether alone or in combination, showed comparable performance (Figure S3). These results suggest that consensus modeling using multiple zero-shot predictors offers a practical strategy to substantially increase the success rate of identifying beneficial mutations, particularly when the mutational search space is moderately large (n ≥ 20).

### Other considerations and concluding remarks

This study highlights several inherent pitfalls of KP predictors that must be considered when applying them to enzyme-directed evolution. First, as previously mentioned, KP data from public databases are often pooled together without aligning the experimental conditions, which can obscure the primary goal of enzyme-directed evolution: identifying sequence-level differences. In this study, we addressed this issue by sorting noisy data into subsets where experimental conditions were consistent across samples, a critical step for accurately evaluating the performance of computational tools.

Second, KP predictors typically use standard loss functions (such as root mean squared error or mean squared error) during training. However, in practice, the relative ranking of mutants is often of greater interest rather than their absolute KP values. Researchers typically aim to predict the relative ranking among mutants to identify variants that outperform the wild type. To address this, we focused on Spearman correlation as a metric to evaluate the ranking performance of computational tools in guiding enzyme-directed evolution.

Additionally, published studies on computationally guided enzyme evolution may be biased toward successful cases, while failures are often underreported. This selective reporting can mislead future researchers, who may overestimate the utility of these tools and continue to invest in computational methods with limited effectiveness. Our study aims to provide a more balanced perspective, revealing both the strengths and limitations of these computational tools.

It is also important to note that, despite our efforts to ensure maximal consistency within subsets, some inconsistencies may still exist in the curated data, such as variations in codon optimization, culture conditions, and experimental protocols. These factors are difficult to resolve without more comprehensive reporting in publications and the development of advanced text processing tools. Another key limitation of our study is that it did not address epistatic interactions, where multiple mutations co-occur and affect each other’s effects. Predicting such epistatic interactions remains a significant challenge, as many predictors (Table 2) are not compatible with these scenarios.

In summary, we evaluated the performance of various commonly used and emerging computational tools for guiding enzyme evolution toward improved catalytic activity. The datasets in this study were carefully curated to ensure consistency, which is a crucial prerequisite for meaningful comparison. Our results highlight the promising potential of large language model-based zero-shot fitness predictors and suggest that mutagenesis data should be sufficiently large to fully leverage the capabilities of computational tools.

## Materials and Methods

### Data Collection from Public Databases

Data were primarily obtained from the BRENDA database (https://www.brenda-enzymes.org/index.php) and the SABIO-RK database (https://sabio.h-its.org/newSearch/index). From BRENDA, entries were collected based on the “Classic view” categories of “Turnover Number,” “*Km* Value,” and “*kcat*/*Km* Value.” For SABIO-RK, data were retrieved by setting the search condition Parametertype to (“*kcat*” OR “*Km*” OR “*kcat*/*Km*”).

### Curation of Independent Datasets

We manually curated 25 additional mutagenesis datasets. The inclusion criteria were:

a. identifiable sequences;
b. small molecule substrate (to ensure compatibility with KP predictors);
c. at least six mutant sequences (including wild type);
d. for *kcat*, *Km*, or *kcat*/*Km* datasts, publication date must be within the latest year (2025) to avoid possible overlap with KP predictor’s training data.

As a result, we managed to curate datasets with a variety of experimental measurements, including relative activity, *kcat*, *Km*, *kcat*/*Km*, and deep mutational scanning (DMS) scores. For the DMS data, the reported fitness scores were averaged across three replicates. Since our downstream evaluation is based on Spearman correlation, which depends on the relative ordering of data points, we first ranked all DMS measurements from low to high, then selected 100 data points at fixed intervals. This strategy reduces the influence of experimental noise and increases the likelihood that adjacent points reflect true biological differences rather than measurement variability, thereby producing a more robust correlation assessment.

### Data Preprocessing

To ensure that the curated datasets were accurate and comparable, strict filtering criteria were applied to the raw entries from BRENDA and SABIO-RK. Entries lacking a UniProt ID, containing multiple UniProt IDs, or missing information on substrate, temperature, pH, or literature reference were removed. Because standardized mutation annotations were not provided in these databases, mutation data were extracted from textual annotations.

We focused on single-point mutations and excluded entries containing the keywords ‘insert’, ‘insertion’, ‘inserted’, ‘delete’, ‘deletion’, ‘deleted’, ‘double’, ‘triple’, ‘multiple’, ‘/’, ‘hybrid’, ‘mixture’, ‘delta’, ‘truncate’, ‘truncated’, or ‘fragment’ (case-insensitive). For SABIO-RK, enzyme types were annotated based on the “Enzyme Variant” field: entries containing “wildtype” were labeled as wildtype enzymes, and those containing “mutate,” “mutated,” “mutation,” “mutant,” “mutagenesis,” or “variant” were labeled as mutated enzymes. For BRENDA, due to the absence of explicit annotations, entries without mutation keywords were considered wildtype.

For entries annotated as mutated enzymes, mutation annotations were retained only if they followed the format of “single amino acid + position number + single amino acid” and involved exactly one mutation. Small datasets (referred to as “subsets”) were constructed as follows: for BRENDA, entries were grouped by EC number, enzyme name, UniProt ID, substrate, organism, reference, temperature, and pH. For SABIO-RK, explicit reaction information was also included in grouping.Only subsets containing one wildtype enzyme and at least nine single-point mutations were retained.

All subsets were then manually validated. Using the UniProt ID, wildtype protein sequences were retrieved from the UniProt database (https://www.uniprot.org/). For each mutation, the amino acid at the specified position was verified against the wildtype sequence. Mutated sequences were generated accordingly. Subsets involving enzymes longer than 1000 amino acids were excluded due to downstream model constraints. For substrate preprocessing, Substrate SMILES representations were obtained as following. InChI identifiers were retrieved under the “Ligands” section and converted to SMILES using RDKit for the BRENDA entries. For SABIO-RK entries, SMILES were obtained from the PubChem database (https://pubchem.ncbi.nlm.nih.gov/).

In total, 441 validated subsets were obtained, including 293 from BRENDA, 121 from SABIO-RK, and 25 from the literature.

### Model Deployment and Execution

Data were analyzed using four categories of models: stability prediction, fitness prediction, enzyme kinetic parameter prediction, and molecular docking affinity prediction. In total, 20 computational models were employed (5 models per category).

#### Stability Prediction Models

Five models were used to assess protein stability changes upon mutation, all using wildtype structures generated by ESMFold. RaSP and ThermoMPNN predict saturation mutagenesis stability scores across all possible substitutions. FoldX provides overall protein stability scores from structural energy calculations. DDGun estimates mutation-specific stability changes (ΔΔG) from the wildtype structure and mutation site. Pythia also evaluates saturation mutagenesis effects using the predicted structure. Together, these models quantify energetic consequences of amino acid substitutions from a structural perspective.

#### Fitness Prediction Models

Mutation effects on protein fitness were evaluated using five models based on sequence and structure representations. GEMME derives fitness scores from multiple sequence alignments produced via MMseqs2. VESPA predicts saturation mutagenesis fitness directly from the wildtype sequence. ESM1V estimates site-specific mutation fitness using transformer-based sequence modeling. SaProt incorporates predicted protein structure and mutation site information. ProSST integrates wildtype sequence, structure, and mutation positions to determine mutation-induced functional changes. These models capture evolutionary constraints and mutation impact on enzyme functionality.

#### Enzyme Kinetic Parameter Prediction Models

Enzymatic parameters were predicted from protein sequences and substrate SMILES using five models. UniKP, EITLEM-Kinetics, and CataPro each predict *kcat*, *Km*, and catalytic efficiency (*kcat*/*Km*). CatPred outputs only *kcat* and *Km*, while DLKcat exclusively predicts *kcat*. All models operate without structural preprocessing, enabling efficient evaluation of catalytic performance from sequence-level inputs.

#### Enzyme-substrate Binding Affinity Prediction and Docking Tools

Five models were employed to predict protein–ligand binding affinity and docking conformations. PSICHIC generates affinity scores using only sequence and SMILES inputs. DynamicBind uses ESMFold-predicted protein structures and SMILES to produce binding scores and docked complexes.

SMINA performs structure-based docking using ESMFold-derived protein PDBQT files and ligand structures converted via RDKit and OpenBabel, with binding sites identified by P2Rank. Boltz-2 predicts affinity and docking complexes using sequence and SMILES representations alone. DSDP follows a structure-based approach similar to SMINA to generate affinity scores and ligand poses. These models collectively evaluate molecular recognition across both sequence-based and structure-based frameworks.

### Adjustment of model prediction directionality

Because different predictors are expected to correlate with experimental measurements in different directions, it is essential to standardize the directionality of the correlations so that higher (positive) correlation values consistently indicate better predictive performance. To enable consistent comparison across models, we therefore established the following working assumptions: a) fitness is assumed to positively correlate with *kcat* and *kcat/Km*, but negatively with *Km*, except in cases where model outputs follow the opposite trend (e.g., VESPA). Using these assumptions, we adjusted the sign of each predictor–measurement correlation so that, after transformation, higher correlation values uniformly represent better performance. The complete mapping of predictors to expected correlation directions is provided in Table S1, and the adjusted results are shown in Figure 3.

### Metric calculation

Spearman correlation, precision, and size-weighted pooled correlation are computed as in Eqn. 1, Eqn. 2, and Eqn. 3.

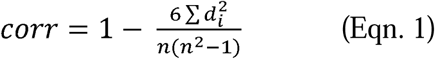

Where d_i_= difference between the ranks of each pair of observations; n=number of pairs.

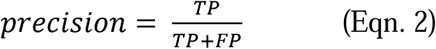

Where TP = True Positives (correctly predicted positive cases); FP = False Positives (incorrectly predicted positive cases).

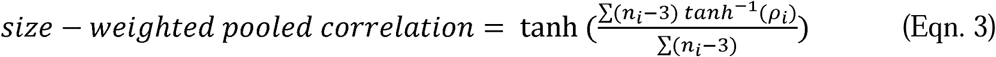

Where n_i_ and p_i_ the number of mutants and correlation for the i-*th* dataset, respectively.

### Wet-lab validation for the PTS mutants

#### Strains and Medium

*Escherichia coli* DH5α was used for plasmid construction and cultured in LB medium supplemented with 100 mg/L ampicillin at 37 °C. The *Saccharomyces cerevisiae* strain yCAN10, previously engineered to enhance the supply of farnesyl pyrophosphate, was used for PTS expression {Luo, 2019 #1299}. Synthetic complete uracil dropout medium (SC-Ura; Coolaber, Beijing, China) was used for yeast selection and cultivation. Yeast extract peptone dextrose (YPD) medium was used for routine yeast culture, and yeast extract peptone galactose (YPG), in which galactose replaced dextrose, was used for fermentation.

#### Mutant recommendation

Saturation mutagenesis of the PTS protein sequence was performed, which were scored and ranked by UniKP (*kcat/Km model*) and ESM-1v—we selected the top 50 mutans recommended by each of these two models. In addition, 50 mutants were randomly selected as the control group. The mutant information (mutation, sequence, and model prediction can be found in Table S2) Primers for these mutants were designed by the Primer3 program {Untergasser, 2012 #1301}.

#### Plasmids and Strains Construction

The patchoulol synthase gene (GenBank ID: AAS86323.1) was synthesized by GenScript (Nanjing, China) with codon optimization for yeast expression. The gene was inserted into plasmid pESC-Ura under the control of the GAL1 promoter and CYC1 terminator using Gibson assembly to generate pESC-Ura-PTS (pGAL1-PTS-tCYC1 expression cassette). Using this plasmid as a template, mutated PTS fragments were amplified by PCR with primers designed to introduce the desired mutations (Table S2). The mutated PTS genes were integrated into the ARS1114a locus of the host strain using CRISPR/Cas9 genome editing {Reider Apel, 2017 #1300}. Briefly, PTS mutational fragments together with ∼1000 bp upstream and downstream homologous arms (amplified from yCAN10 genomic DNA) and the Cas9/sgRNA plasmid p426_Cas9_gRNA-ARS1114a (Addgene #87397) were transformed into the host strain using the lithium acetate method. Transformants were selected on SD-Ura plates and verified by colony PCR and sequencing. All primer sequences used in this study are listed in Table S3.

#### Fermentation of PTS Mutant Strains

Single colonies of mutant strains were inoculated into 96-deep-well plates containing 500 µL YPD per well and cultured at 30 °C and 800 rpm for 18 h to obtain primary seed cultures. Cultures were then transferred to 24-deep-well plates containing 2 mL YPG per well at a 2% inoculum and incubated under the same conditions. All strains were cultured in three biological replicates. After 72 h of fermentation, 500 µL of broth was collected for patchoulol quantification.

#### Extraction and detection methods

Patchoulol was extracted with 200 µL *n*-hexane using bead beating (70 Hz; 90 s on, 30 s off; 10 cycles), followed by centrifugation at 12,000 ×g for 5 min. The upper organic phase was collected for analysis. Patchoulol was quantified by GC-MS (8890-7010B, Agilent, USA) equipped with an HP-5 column (30 m × 250 µm × 0.25 µm, Agilent). The oven temperature program was: 150 °C for 1 min, ramp to 180 °C at 20 °C/min, then to 300 °C at 80 °C/min and held for 4 min. A 1 µL sample was injected with helium as the carrier gas. Selected ion monitoring (SIM) was used for quantification with *m/z* 137.9, 160.9, and 222. The mutants’ patchoulol yield relative to wildtype can be found in Table S2.

## Data and code availability

Data in this study, including curated subsets, experimental conditions, experimental measurement, and pre-computed predictions, are deposited in our GitHub repository. (https://github.com/zishuozeng/EnzyArena).

## Ethical statement

Not applicable.

## Acknowledgements

We would like to thank all researchers who made the data and tools relevant to this study available.

## Authorship contribution

Z.Z. and X.L. conceived the ideas, analyzed the data, and wrote the manuscript; J.J. collected the data and ran the predictions; R.X. performed the wet-lab experiments; X.L. revised the manuscript.

## References

(1) Radley, E.; Davidson, J.; Foster, J.; Obexer, R.; Bell, E. L.; Green, A. P. Engineering enzymes for environmental sustainability. Angewandte Chemie International Edition 2023, 62 (52), e202309305.

(2) Katsimpouras, C.; Stephanopoulos, G. Enzymes in biotechnology: Critical platform technologies for bioprocess development. Current opinion in biotechnology 2021, 69, 91–102.

(3) Markus, B.; C, G. C.; Andreas, K.; Arkadij, K.; Stefan, L.; Gustav, O.; Elina, S.; Radka, S. Accelerating biocatalysis discovery with machine learning: a paradigm shift in enzyme engineering, discovery, and design. ACS catalysis 2023, 13 (21), 14454–14469.

(4) Meier, J.; Rao, R.; Verkuil, R.; Liu, J.; Sercu, T.; Rives, A. Language models enable zero-shot prediction of the effects of mutations on protein function. Advances in neural information processing systems 2021, 34, 29287–29303.

(5) Elnaggar, A.; Heinzinger, M.; Dallago, C.; Rihawi, G.; Wang, Y.; Jones, L.; Gibbs, T.; Feher, T.; Angerer, C.; Steinegger, M. ProtTrans: towards cracking the language of Life’s code through self-supervised deep learning and high performance computing. arXiv preprint arXiv:2007.06225 2020.

(6) Su, J.; Han, C.; Zhou, Y.; Shan, J.; Zhou, X.; Yuan, F. Saprot: Protein language modeling with structure-aware vocabulary. BioRxiv 2023, 2023.2010. 2001.560349.

(7) Li, M.; Tan, Y.; Ma, X.; Zhong, B.; Yu, H.; Zhou, Z.; Ouyang, W.; Zhou, B.; Tan, P.; Hong, L. Prosst: Protein language modeling with quantized structure and disentangled attention. Advances in Neural Information Processing Systems 2024, 37, 35700–35726.

(8) Li, F.; Yuan, L.; Lu, H.; Li, G.; Chen, Y.; Engqvist, M. K.; Kerkhoven, E. J.; Nielsen, J. Deep learning-based k cat prediction enables improved enzyme-constrained model reconstruction. Nature Catalysis 2022, 5 (8), 662–672.

(9) Yu, H.; Deng, H.; He, J.; Keasling, J. D.; Luo, X. UniKP: a unified framework for the prediction of enzyme kinetic parameters. Nature communications 2023, 14 (1), 8211.

(10) Wang, J.; Yang, Z.; Chen, C.; Yao, G.; Wan, X.; Bao, S.; Ding, J.; Wang, L.; Jiang, H. MPEK: a multitask deep learning framework based on pretrained language models for enzymatic reaction kinetic parameters prediction. Briefings in Bioinformatics 2024, 25 (5), bbae387.

(11) Wang, Z.; Xie, D.; Wu, D.; Luo, X.; Wang, S.; Li, Y.; Yang, Y.; Li, W.; Zheng, L. Robust enzyme discovery and engineering with deep learning using CataPro. Nature communications 2025, 16 (1), 2736.

(12) Boorla, V. S.; Maranas, C. D. CatPred: a comprehensive framework for deep learning in vitro enzyme kinetic parameters. Nature communications 2025, 16 (1), 2072.

(13) Koes, D. R.; Baumgartner, M. P.; Camacho, C. J. Lessons learned in empirical scoring with smina from the CSAR 2011 benchmarking exercise. Journal of chemical information and modeling 2013, 53 (8), 1893–1904.

(14) Lu, W.; Zhang, J.; Huang, W.; Zhang, Z.; Jia, X.; Wang, Z.; Shi, L.; Li, C.; Wolynes, P. G.; Zheng, S. DynamicBind: predicting ligand-specific protein-ligand complex structure with a deep equivariant generative model. Nature Communications 2024, 15 (1), 1071.

(15) Koh, H. Y.; Nguyen, A. T.; Pan, S.; May, L. T.; Webb, G. I. Physicochemical graph neural network for learning protein–ligand interaction fingerprints from sequence data. Nature Machine Intelligence 2024, 6 (6), 673–687.

(16) Huang, Y.; Zhang, H.; Jiang, S.; Yue, D.; Lin, X.; Zhang, J.; Gao, Y. Q. Dsdp: A blind docking strategy accelerated by gpus. Journal of chemical information and modeling 2023, 63 (14), 4355–4363.

(17) Passaro, S.; Corso, G.; Wohlwend, J.; Reveiz, M.; Thaler, S.; Somnath, V. R.; Getz, N.; Portnoi, T.; Roy, J.; Stark, H. Boltz-2: Towards accurate and efficient binding affinity prediction. BioRxiv 2025.

(18) Delgado, J.; Radusky, L. G.; Cianferoni, D.; Serrano, L. FoldX 5.0: working with RNA, small molecules and a new graphical interface. Bioinformatics 2019, 35 (20), 4168–4169.

(19) Blaabjerg, L. M.; Kassem, M. M.; Good, L. L.; Jonsson, N.; Cagiada, M.; Johansson, K. E.; Boomsma, W.; Stein, A.; Lindorff-Larsen, K. Rapid protein stability prediction using deep learning representations. Elife 2023, 12, e82593.

(20) Dieckhaus, H.; Brocidiacono, M.; Randolph, N. Z.; Kuhlman, B. Transfer learning to leverage larger datasets for improved prediction of protein stability changes. Proceedings of the national academy of sciences 2024, 121 (6), e2314853121.

(21) Montanucci, L.; Capriotti, E.; Frank, Y.; Ben-Tal, N.; Fariselli, P. DDGun: an untrained method for the prediction of protein stability changes upon single and multiple point variations. BMC bioinformatics 2019, 20, 1–10.

(22) Sun, J.; Zhu, T.; Cui, Y.; Wu, B. Structure-based self-supervised learning enables ultrafast protein stability prediction upon mutation. The Innovation 2025, 6 (1).

(23) Cui, Y.; Sun, J.; Wu, B. Computational enzyme redesign: large jumps in function. Trends in Chemistry 2022, 4 (5), 409–419.

(24) Mangul, S.; Martin, L. S.; Hill, B. L.; Lam, A. K.-M.; Distler, M. G.; Zelikovsky, A.; Eskin, E.; Flint, J. Systematic benchmarking of omics computational tools. Nature communications 2019, 10 (1), 1393. Camacho, D. M.; Collins, K. M.; Powers, R. K.; Costello, J. C.; Collins, J. J. Next-generation machine learning for biological networks. Cell 2018, 173 (7), 1581-1592. Pleiss, J. r. Modeling Enzyme Kinetics: Current Challenges and Future Perspectives for Biocatalysis. Biochemistry 2024, 63 (20), 2533-2541.

(25) Chang, A.; Jeske, L.; Ulbrich, S.; Hofmann, J.; Koblitz, J.; Schomburg, I.; Neumann-Schaal, M.; Jahn, D.; Schomburg, D. BRENDA, the ELIXIR core data resource in 2021: new developments and updates. Nucleic acids research 2021, 49 (D1), D498–D508.

(26) Wittig, U.; Rey, M.; Weidemann, A.; Kania, R.; Müller, W. SABIO-RK: an updated resource for manually curated biochemical reaction kinetics. Nucleic acids research 2018, 46 (D1), D656–D660.

(27) Kapoor, S.; Narayanan, A. Leakage and the reproducibility crisis in machine-learning-based science. Patterns 2023, 4 (9).

(28) Bernett, J.; Blumenthal, D. B.; Grimm, D. G.; Haselbeck, F.; Joeres, R.; Kalinina, O. V.; List, M. Guiding questions to avoid data leakage in biological machine learning applications. Nature Methods 2024, 21 (8), 1444–1453.

(29) Graber, D.; Stockinger, P.; Meyer, F.; Mishra, S.; Horn, C.; Buller, R. Resolving data bias improves generalization in binding affinity prediction. Nature Machine Intelligence 2025, 1–13.

(30) Notin, P.; Kollasch, A.; Ritter, D.; Van Niekerk, L.; Paul, S.; Spinner, H.; Rollins, N.; Shaw, A.; Orenbuch, R.; Weitzman, R. Proteingym: Large-scale benchmarks for protein fitness prediction and design. Advances in Neural Information Processing Systems 2023, 36, 64331–64379.

(31) Siddiqui, K. S. Defying the activity–stability trade-off in enzymes: taking advantage of entropy to enhance activity and thermostability. Critical reviews in biotechnology 2017, 37 (3), 309–322.

(32) Vanella, R.; Küng, C.; Schoepfer, A. A.; Doffini, V.; Ren, J.; Nash, M. A. Understanding activity-stability tradeoffs in biocatalysts by enzyme proximity sequencing. Nature Communications 2024, 15 (1), 1807.

(33) Zheng, N.; Cai, Y.; Zhang, Z.; Zhou, H.; Deng, Y.; Du, S.; Tu, M.; Fang, W.; Xia, X. Tailoring industrial enzymes for thermostability and activity evolution by the machine learning-based iCASE strategy. Nature Communications 2025, 16 (1), 604. Jiang, F.; Li, M.; Dong, J.; Yu, Y.; Sun, X.; Wu, B.; Huang, J.; Kang, L.; Pei, Y.; Zhang, L. A general temperature-guided language model to design proteins of enhanced stability and activity. Science Advances 2024, 10 (48), eadr2641. Teufl, M.; Zajc, C. U.; Traxlmayr, M. W. Engineering strategies to overcome the stability–function trade-off in proteins. ACS *Synthetic Biology* 2022, 11 (3), 1030-1039.

(34) Yu, Y.; Rué Casamajo, A.; Finnigan, W.; Schnepel, C.; Barker, R.; Morrill, C.; Heath, R. S.; De Maria, L.; Turner, N. J.; Scrutton, N. S. Structure-based design of small imine reductase panels for target substrates. ACS catalysis 2023, 13 (18), 12310–12321. Qian, S.; Clomburg, J. M.; Gonzalez, R. Engineering Escherichia coli as a platform for the in vivo synthesis of prenylated aromatics. Biotechnology and Bioengineering 2019, 116 (5), 1116-1127. Mai, P.; Zocher, G.; Stehle, T.; Li, S.-M. Structure-based protein engineering enables prenyl donor switching of a fungal aromatic prenyltransferase. Organic & Biomolecular Chemistry 2018, 16 (40), 7461-7469. Hua, L.; Qianqian, B.; Jianfeng, Z.; Yinbiao, X.; Shengyu, Y.; Weishi, X.; Yang, S.; Yupeng, L. Directed evolution engineering to improve activity of glucose dehydrogenase by increasing pocket hydrophobicity. *Frontiers in Microbiology* 2022, 13, 1044226.

(35) Laine, E.; Karami, Y.; Carbone, A. GEMME: a simple and fast global epistatic model predicting mutational effects. Molecular biology and evolution 2019, 36 (11), 2604–2619.

(36) Zhang, Q.; Chen, W.; Qin, M.; Wang, Y.; Pu, Z.; Ding, K.; Liu, Y.; Zhang, Q.; Li, D.; Li, X. Integrating protein language models and automatic biofoundry for enhanced protein evolution. Nature Communications 2025, 16 (1), 1553.

(37) Cornish-Bowden, A. Fundamentals of enzyme kinetics; John Wiley & Sons, 2013.

(38) Im, W.; Lee, M. S.; Brooks III, C. L. Generalized born model with a simple smoothing function. Journal of computational chemistry 2003, 24 (14), 1691–1702.

(39) Dieckhaus, H.; Kuhlman, B. Protein stability models fail to capture epistatic interactions of double point mutations. Protein Science 2025, 34 (1), e70003.

(40) James, G.; Witten, D.; Hastie, T.; Tibshirani, R. An introduction to statistical learning; Springer, 2013.

(41) Fisher, R. A. On the’ probable error’ of a coefficient of correlation deduced from a small sample. Metron 1921, 1, 3–32.

(42) Wrenbeck, E. E.; Azouz, L. R.; Whitehead, T. A. Single-mutation fitness landscapes for an enzyme on multiple substrates reveal specificity is globally encoded. Nature communications 2017, 8 (1), 15695.

(43) Muir, D. F.; Asper, G. P.; Notin, P.; Posner, J. A.; Marks, D. S.; Keiser, M. J.; Pinney, M. M. Evolutionary-scale enzymology enables exploration of a rugged catalytic landscape. Science 2025, 388 (6752), eadu1058.

(44) Luo, S.; Hou, Y.; Hu, S.-Q. Enhancement of preference, catalytic activity and thermostability of polyphenol oxidase from Rosa Chinensis by semi-rational engineering. Molecular Catalysis 2024, 559, 114059.

(45) Marquet, C.; Heinzinger, M.; Olenyi, T.; Dallago, C.; Erckert, K.; Bernhofer, M.; Nechaev, D.; Rost, B. Embeddings from protein language models predict conservation and variant effects. Human genetics 2022, 141 (10), 1629–1647.

(46) Shen, X.; Cui, Z.; Long, J.; Zhang, S.; Chen, B.; Tan, T. EITLEM-Kinetics: A deep-learning framework for kinetic parameter prediction of mutant enzymes. Chem Catalysis 2024, 4 (9).

