## Supplementary Information for "Benchmarking and Experimental Validation of Machine Learning Strategies for Enzyme Engineering"

**Detailed descriptions of the selected predictors**

**Protein-ligand binding affinity prediction**: Protein–ligand binding affinity prediction is a widely adopted strategy for assessing enzyme–substrate interactions ^1,2^. For this strategy, we selected SMINA ^3^, DynamicBind ^4^, PSICHIC ^5^, DSDP ^6^, and Boltz-2 ^7^. SMINA is an open-source, physics-based molecular protein-ligand binding affinity prediction program designed to provide flexible and accurate predictions of small molecule binding in protein binding sites. It is a fork of the widely used AutoDock Vina ^8^ and offers customizable scoring functions. SMINA has been reported to outperform other open-source, physics-based tools in protein-ligand binding affinity prediction ^9^. DynamicBind and PSICHIC, in contrast, are deep learning frameworks, making them fundamentally different from traditional tools like SMINA. DynamicBind focuses on modeling the dynamic nature of proteins, emphasizing the ability to capture ligand-induced conformational changes and recover ligand-specific protein structures. It utilizes equivariant geometric diffusion networks to construct smooth energy landscapes for efficient transitions between equilibrium states. PSICHIC, on the other hand, takes a sequence-based approach, bypassing structural or dynamic data to directly decode protein-ligand interaction fingerprints from sequence information, making it structure-agnostic. DSDP combines traditional docking and machine learning to enable fast blind docking by predicting the initial binding position. Boltz-2 is a recent AI foundation model that predicts both the structure and binding affinity from interactions between small molecules/biomacromolecules and proteins.

**Stability prediction**: Protein stability prediction plays a complementary role in enzyme design, as mutations that enhance catalytic activity may simultaneously reduce structural stability, highlighting an intrinsic trade-off between the two properties ^10,11^. We selected FoldX ^12^, RaSP ^13^, ThermoMPNN ^14^, DDGun ^15^, and Pythia ^16^ for this strategy. FoldX is a well-established and commonly used tool that evaluates the effects of mutations on protein stability using empirical force fields and energy functions. It has been applied to various fields, including nuclease design ^17^, lyase catalytic activity improvement ^18^, and deleterious mutant identification ^19^. RaSP and ThermoMPNN are newer deep learning-based predictors that have been shown to outperform FoldX in stability prediction ^14,20^. RaSP uses a deep learning architecture for fast, scalable prediction of saturated mutagenesis outcomes. ThermoMPNN, based on transfer learning, utilizes embeddings derived from ProteinMPNN ^21^ as feature inputs. DDGun is a computational model for predicting the protein stability changes upon mutations using predefined rules derived from sequence, structure, and evolutionary conservation. A recent benchmark study highlighted the superior performances of RaSP, ThermoMPNN, and DDGun among 27 models for protein stability prediction ^20^. Pythia is a recent model for fast mutations-induced protein stability prediction based on a self-supervised graph neural network.

**Zero-shot fitness prediction**: Zero-shot fitness prediction estimates mutational effects without task-specific training, relying instead on generalizable representations learned from large-scale protein data. For this strategy, we selected GEMME ^22^, VESPA ^23^, ESM-1v ^24^, SaProt ^25^ and ProSST ^26^ to represent evolutionary information-driven and large language model (LLM)-based approaches for protein fitness prediction. GEMME (Global Epistatic Model for Mutational Effects) is a computational model that integrates conservation, evolutionary fit, and site-independent frequencies for fitness prediction. VESPA and ESM-1v are part of two major protein LLM projects—ProtTrans ^27^ and Evolutionary Scale Modeling (ESM) [<https://github.com/facebookresearch/esm>], respectively. ProtTrans includes several transformer-based LLM models, such as ProtT5, ProtBert, and ProAlbert, which are mainly used for protein embeddings in transfer learning applications. VESPA, trained on datasets of single amino acid variant (SAV) effects, is one such transfer learning application. The ESM series includes models like ESM2 ^28^ (for embeddings and general purposes), ESMFold ^28^ (for protein structure prediction), ESM-1v ^24^ (for variant effect prediction), and ESM-IF1 ^29^ (for inverse folding). Both SaProt and ProSST seek to incorporate protein structural information into training of language model. By doing so, SaProt adopts a new type of tokens that carry both sequence and structural information; while ProSST equips with a structure quantization module that converts local protein structures into discrete tokens. SaProt and ProSST have temporarily topped the ProteinGym ^30^ benchmark and ProSST is still the third place as of today.

Notably, the zero-shot predictors selected in this study differ in the way they interpret and score their predictions. For ESM-1v, SaProt, and ProSST, the scores reflect how well a variant "fits" within a protein context. A high positive score indicates that the variant is likely to be compatible within the protein sequence and/or structure, suggesting a potential for enhanced function or stability. Conversely, a negative score implies that the variant may disrupt the sequence, potentially reducing function or destabilizing the structure. In contrast, GEMME's scores indicate the deleterious nature of a variant when positive and low/negative scores suggest minimal or neutral effects. VESPA outputs a probability score for a binary label—effect or neutral—where the term "effect" encompasses both beneficial and deleterious outcomes. From an evolutionary perspective, a variant with an "effect" label is more likely to be deleterious rather than beneficial ^31^.

**Enzyme kinetic parameter (KP) prediction**: Enzyme kinetic parameter (KP) prediction targets key catalytic descriptors—turnover number (*kcat*), Michaelis constant (*Km*), and catalytic efficiency (*kcat*/*Km*)—which are critical for characterizing and optimizing enzyme function. Recently many models have been developed to directly predict these KPs, from which we selected DLKcat^32^, UniKP ^33^, EITLEM-kinetics ^34^, CataPro ^35^ and CatPred ^36^. These predictors take enzyme sequences and small molecule substrate as input and predict the KP, such as *kcat*, *Km*, and/or *kcat*/*Km*. The training data of these predictors are derived primarily from BRENDA and SABIO-RK. Notably, EITLEM-kinetics is trained exclusively on mutant datasets and is designed to prioritize enzyme mutants that improve catalytic activity. In contrast, other predictors incorporate both mutant and wildtype data in their training. DLKcat used a n-gram strategy for protein feature extraction; while other four predictors used pre-trained protein language models (ProtT5, ESM2, or ESM-1v) instead.

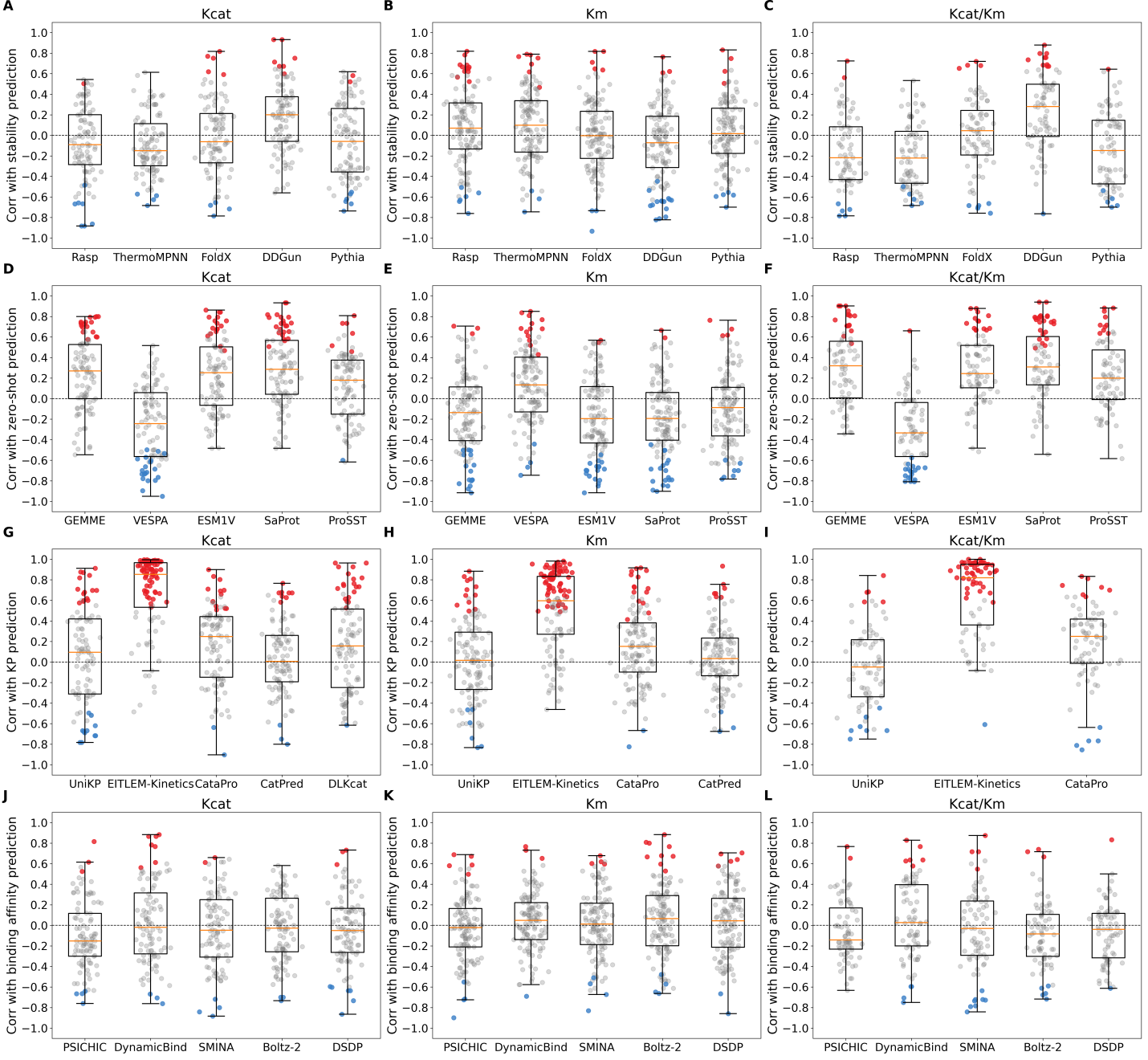

**Figure S1. Performances overview of computational tools on subsets derived from BRENDA.** Performance is measured by Spearman correlation. In each panel, red, blue, and grey dots indicate significantly (p-value < 0.05) positive, significantly negative, and non-significant Spearman correlations, respectively.

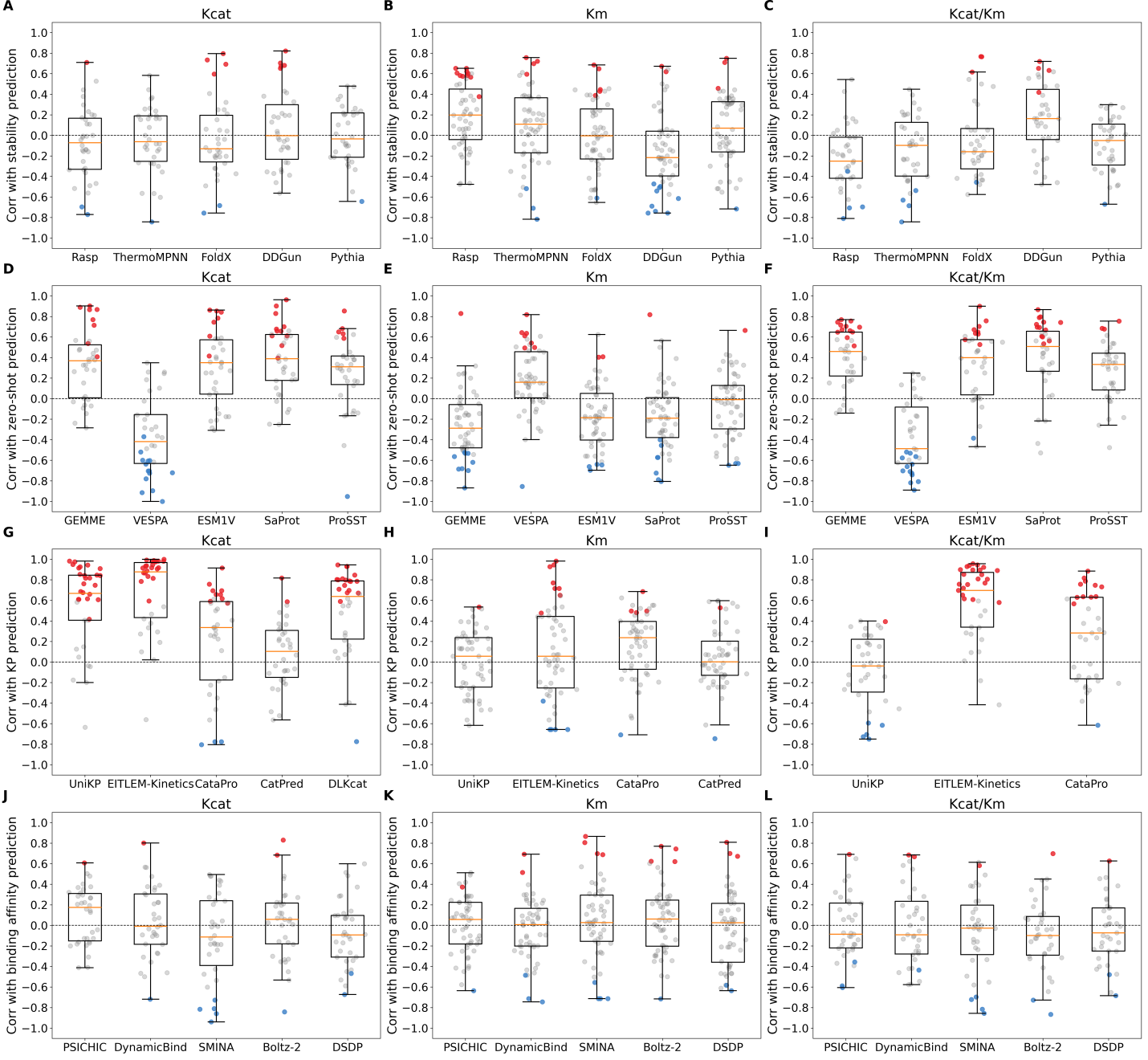

**Figure S2. Performances overview of computational tools on subsets derived from SABIO-RK.** Performance is measured by Spearman correlation. In each panel, red, blue, and grey dots indicate significantly (p-value < 0.05) positive, significantly negative, and non-significant Spearman correlations, respectively.

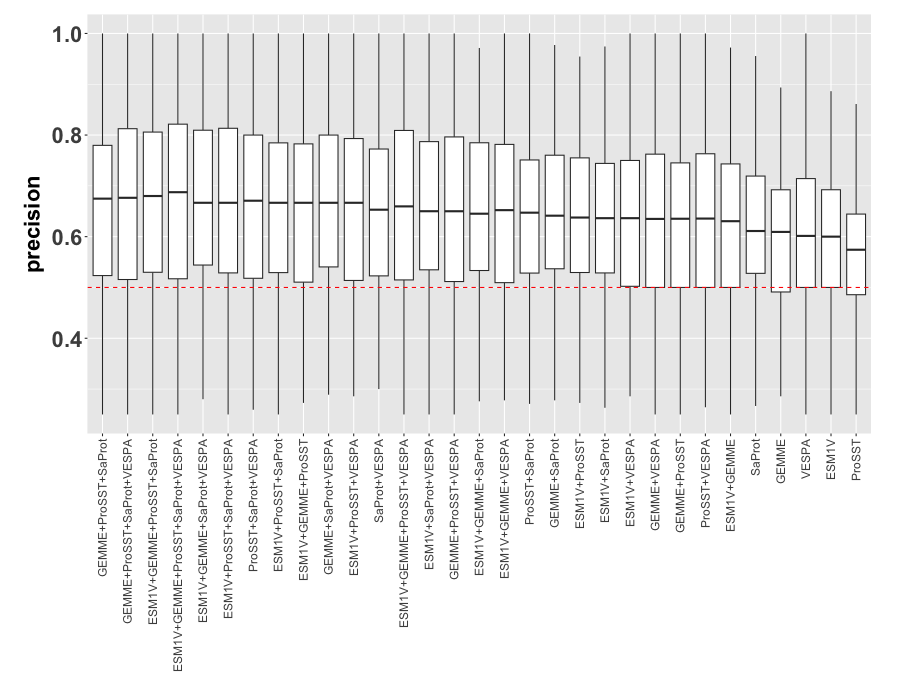

**Figure S3. Combination of zero-shot predictors does not improve precision in identifying beneficial enzyme mutants when sample size is small.** (A) Precision of all 31 combinations of five zero-shot predictors, evaluated using exhaustive pairwise comparisons across datasets without setting a sample size cutoff—as a result, most datasets are of small sample size. Each box plot represents the distribution of precision values across datasets, with the dashed red line indicating random baseline performance. No significant difference is observed between any two combo or single predictions (p > 0.05, Wilcoxon rank-sum test).

**Table S1. Adjustment of model prediction’s directionality.**

| Category | Model | Output | Assgined direction | | |
| --- | --- | --- | --- | --- | --- |
|  |  |  | Kcat Corr | Km Corr | Kcat/Km Corr |
| Stability | Pythia | ΔΔG | + | - | + |
|  | ThermoMPNN | ΔΔG | + | - | + |
|  | DDGun* | ΔΔG | - | + | - |
|  | Rasp | ΔΔG | + | - | + |
|  | FoldX | ΔG | + | - | + |
| Kinetic Parameters | DLKcat | Kcat | + | / | / |
|  | UniKP | Kcat, Km, Kcat/Km | + | + | + |
|  | CataPro | Kcat, Km, Kcat/Km | + | + | + |
|  | CatPred | Kcat, Km | + | + | / |
|  | EITLEM-Kinetic Parameters | Kcat, Km, Kcat/Km | + | + | + |
| Fitness | SaProt | Variant Effect Score | + | - | + |
|  | ESM1V | Variant Effect Score | + | - | + |
|  | ProSST | Variant Effect Score | + | - | + |
|  | GEMME | Variant Effect Score | + | - | + |
|  | VESPA | Variant Effect Score | - | + | - |
| Binding Affinity | PSICHIC | Binding Affinity | + | - | + |
|  | DSDP | Binding Affinity | - | + | - |
|  | SMINA | Binding Affinity | - | + | - |
|  | DynamicBind | Binding Affinity | + | - | + |
|  | Boltz-2 | Binding Affinity | - | + | - |

*DDGun’s GitHub repository (<https://github.com/biofold/ddgun>) indicates that higher predictions correspond to higher stability.

1	Xiong, W., Liu, B., Shen, Y., Jing, K. & Savage, T. R. Protein engineering design from directed evolution to de novo synthesis. *Biochemical Engineering Journal* **174**, 108096 (2021).

2	Marshall, L. R., Bhattacharya, S. & Korendovych, I. V. Fishing for Catalysis: Experimental Approaches to Narrowing Search Space in Directed Evolution of Enzymes. *JACS Au* **3**, 2402-2412 (2023).

3	Koes, D. R., Baumgartner, M. P. & Camacho, C. J. Lessons learned in empirical scoring with smina from the CSAR 2011 benchmarking exercise. *Journal of chemical information and modeling* **53**, 1893-1904 (2013).

4	Lu, W. *et al.* DynamicBind: predicting ligand-specific protein-ligand complex structure with a deep equivariant generative model. *Nature Communications* **15**, 1071 (2024).

5	Koh, H. Y., Nguyen, A. T., Pan, S., May, L. T. & Webb, G. I. Physicochemical graph neural network for learning protein–ligand interaction fingerprints from sequence data. *Nature Machine Intelligence* **6**, 673-687 (2024).

6	Huang, Y. *et al.* Dsdp: A blind docking strategy accelerated by gpus. *Journal of chemical information and modeling* **63**, 4355-4363 (2023).

7	Passaro, S. *et al.* Boltz-2: Towards accurate and efficient binding affinity prediction. *BioRxiv* (2025).

8	Trott, O. & Olson, A. J. AutoDock Vina: improving the speed and accuracy of docking with a new scoring function, efficient optimization, and multithreading. *Journal of computational chemistry* **31**, 455-461 (2010).

9	Masters, L., Eagon, S. & Heying, M. Evaluation of consensus scoring methods for AutoDock Vina, smina and idock. *Journal of Molecular Graphics and Modelling* **96**, 107532 (2020).

10	Siddiqui, K. S. Defying the activity–stability trade-off in enzymes: taking advantage of entropy to enhance activity and thermostability. *Critical reviews in biotechnology* **37**, 309-322 (2017).

11	Vanella, R. *et al.* Understanding activity-stability tradeoffs in biocatalysts by enzyme proximity sequencing. *Nature Communications* **15**, 1807 (2024).

12	Delgado, J., Radusky, L. G., Cianferoni, D. & Serrano, L. FoldX 5.0: working with RNA, small molecules and a new graphical interface. *Bioinformatics* **35**, 4168-4169 (2019).

13	Blaabjerg, L. M. *et al.* Rapid protein stability prediction using deep learning representations. *Elife* **12**, e82593 (2023).

14	Dieckhaus, H., Brocidiacono, M., Randolph, N. Z. & Kuhlman, B. Transfer learning to leverage larger datasets for improved prediction of protein stability changes. *Proceedings of the national academy of sciences* **121**, e2314853121 (2024).

15	Montanucci, L., Capriotti, E., Frank, Y., Ben-Tal, N. & Fariselli, P. DDGun: an untrained method for the prediction of protein stability changes upon single and multiple point variations. *BMC bioinformatics* **20**, 1-10 (2019).

16	Sun, J., Zhu, T., Cui, Y. & Wu, B. Structure-based self-supervised learning enables ultrafast protein stability prediction upon mutation. *The Innovation* **6** (2025).

17	He, Z., Mei, G., Zhao, C. & Chen, Y. Potential application of FoldX force field based protein modeling in zinc finger nucleases design. *Science China Life Sciences* **54**, 442-449 (2011).

18	Huang, A. *et al.* Improving the thermal stability and catalytic activity of ulvan lyase by the combination of FoldX and KnowVolution campaign. *International Journal of Biological Macromolecules* **257**, 128577 (2024).

19	Gerasimavicius, L., Livesey, B. J. & Marsh, J. A. Loss-of-function, gain-of-function and dominant-negative mutations have profoundly different effects on protein structure. *Nature communications* **13**, 3895 (2022).

20	Zheng, F., Liu, Y., Yang, Y., Wen, Y. & Li, M. Assessing computational tools for predicting protein stability changes upon missense mutations using a new dataset. *Protein Science* **33**, e4861 (2024).

21	Dauparas, J. *et al.* Robust deep learning–based protein sequence design using ProteinMPNN. *Science* **378**, 49-56 (2022).

22	Laine, E., Karami, Y. & Carbone, A. GEMME: a simple and fast global epistatic model predicting mutational effects. *Molecular biology and evolution* **36**, 2604-2619 (2019).

23	Marquet, C. *et al.* Embeddings from protein language models predict conservation and variant effects. *Human genetics* **141**, 1629-1647 (2022).

24	Meier, J. *et al.* Language models enable zero-shot prediction of the effects of mutations on protein function. *Advances in neural information processing systems* **34**, 29287-29303 (2021).

25	Su, J. *et al.* Saprot: Protein language modeling with structure-aware vocabulary. *BioRxiv*, 2023.2010. 2001.560349 (2023).

26	Li, M. *et al.* Prosst: Protein language modeling with quantized structure and disentangled attention. *Advances in Neural Information Processing Systems* **37**, 35700-35726 (2024).

27	Elnaggar, A. *et al.* ProtTrans: towards cracking the language of Life's code through self-supervised deep learning and high performance computing. *arXiv preprint arXiv:2007.06225* (2020).

28	Lin, Z. *et al.* Evolutionary-scale prediction of atomic-level protein structure with a language model. *Science* **379**, 1123-1130 (2023).

29	Hsu, C. *et al.* in *International conference on machine learning.* 8946-8970 (PMLR).

30	Notin, P. *et al.* Proteingym: Large-scale benchmarks for protein fitness prediction and design. *Advances in Neural Information Processing Systems* **36**, 64331-64379 (2023).

31	Zeng, Z., Aptekmann, A. A. & Bromberg, Y. Decoding the effects of synonymous variants. *Nucleic acids research* **49**, 12673-12691 (2021).

32	Li, F. *et al.* Deep learning-based k cat prediction enables improved enzyme-constrained model reconstruction. *Nature Catalysis* **5**, 662-672 (2022).

33	Yu, H., Deng, H., He, J., Keasling, J. D. & Luo, X. UniKP: a unified framework for the prediction of enzyme kinetic parameters. *Nature communications* **14**, 8211 (2023).

34	Shen, X. *et al.* EITLEM-Kinetics: A deep-learning framework for kinetic parameter prediction of mutant enzymes. *Chem Catalysis* **4** (2024).

35	Wang, Z. *et al.* Robust enzyme discovery and engineering with deep learning using CataPro. *Nature communications* **16**, 2736 (2025).

36	Boorla, V. S. & Maranas, C. D. CatPred: a comprehensive framework for deep learning in vitro enzyme kinetic parameters. *Nature communications* **16**, 2072 (2025).
