## Supplementary material for "Benchmarking and Experimental Validation of Machine Learning Strategies for Enzyme Engineering": Table S3

**Table S3 Primers used in this study**

| Primers | Primer sequence |
| --- | --- |
| PTS-pGAL1-F | TCAAGGAGAAAAAACATGGAGTTGTATGCCCAA |
| PTS-tCYC1-R | TCGGTTAGAGCGGATTTAATATGGAACAGGGTG |
| pESC-Ura backbone-F | ATCCGCTCTAACCGAAAAGGAAGGA |
| pESC-Ura backbone-R | GTTTTTTCTCCTTGACGTTA |
| pGAL1-1114a-F | ATTACTGAAAGTTCCAAAGAGAAGGGATAATAGTACAAACTTACATAGCGT |
| Tcyc1-1114a-R | TGAGAAGGTTTTGGGACGCTCGAAGAGGACAGTCAATAGCATCAT |
| PTS-V193Y-F | CCGCGAAACAAtatCACAACGCATTGAATGAGTTCTCT |
| PTS-V193S-F | CGCGAAACAAtctCACAACGCATTGAATGAGTTCTCT |
| PTS-H462F-F | ACGAGAGtttGTTCGCACTGCAGTAGAATGCTA |
| PTS-V193W-F | GAAACAAtggCACAACGCATTGAATGAGTTCTCT |
| PTS-V193R-F | CGCGAAACAAagaCACAACGCATTGAATGAGTTCTCT |
| PTS-D187P-F | TTTGAACccaCCAACCGCGAAACAAGTCC |
| PTS-G499F-F | GAGtttTTCCTCAGACCAACAGAATTTCC |
| PTS-V288M-F | ATTCGatgGCTAGGATGATTTTAGCAAAAGGG |
| PTS-Q488L-F | CAACttgATGGAGTCAGCATGGAAGGACA |
| PTS-L178P-F | GAACTACccaGAATCAGTCTATGCAACTTTGAACG |
| PTS-A183H-F | catACTTTGAACGATCCAACCGCG |
| PTS-L56Y-F | AGAGAGtatAAGGAAGCATCAGACAACTACATGC |
| PTS-A241F-F | AtttTTGCACAGAAGGGAGCTGAGTG |
| PTS-L56W-F | AAGAGAGtggAAGGAAGCATCAGACAACTACATGC |
| PTS-H194K-F | CCGCGAAACAAGTCaaaAACGCATTGAATGAGTTCTCTTTTC |
| PTS-E46P-F | AGGCTccaGAGCTGAAAGTGGAGCTGAAAAG |
| PTS-A241I-F | AattTTGCACAGAAGGGAGCTGAGTG |
| PTS-H225Y-F | CATCTCATtatAAAGGCTTGCTCAAACTTGCTAA |
| PTS-G146L-F | TGAGGATttgGCGGTTGCAGTCCTTGAATT |
| PTS-P362F-F | AGGCGCAtttTATCGAGCCTACTATGGAAAAGAAGC |
| PTS-V83W-F | TtggGAAGATGTTGATGAAGCTTTGAAGA |
| PTS-Q488M-F | CAACatgATGGAGTCAGCATGGAAGGACA |
| PTS-Y367L-F | AGCCTACttgGGAAAAGAAGCCATGAAATACGC |
| PTS-H225K-F | CATCTCATaaaAAAGGCTTGCTCAAACTTGCTAA |
| PTS-V467L-F | CACTGCAttgGAATGCTACATGGAAGAGCACAAA |
| PTS-G499C-F | TGAGtgtTTCCTCAGACCAACAGAATTTCC |
| PTS-A309P-F | GTGTATGATccaTATGGTACTTTTGAGGAATTACAAATGT |
| PTS-K478W-F | AAAGTGGGGtggCAAGAGGCCGTGTCTGAATTCT |
| PTS-P188M-F | GAACGATatgACCGCGAAACAAGTCCACAA |
| PTS-M372S-F | GAAGCCtctAAATACGCCGCGAGAGCTT |
| PTS-R267P-F | CATTCGTTccaGATCGATTGGTGGAGTCCTACTTC |
| PTS-V193Q-F | CGCGAAACAAcaaCACAACGCATTGAATGAGTTCTCT |
| PTS-V10F-F | TGTTGGAtttGGTGCTGCTTCTCGTCCTCTT |
| PTS-S14V-F | CTGCTgttCGTCCTCTTGCGAATTTTCATC |
| PTS-A156Y-F | CTTCGAAtatACGCATCTCAGAGTCCATGGA |
| PTS-P335W-F | AACTTtggGATTACATGAAAATAGTATACAAGGCCC |
| PTS-R204L-F | TTTCGAttgGGATTGCCACGCGTGGAA |
| PTS-I215A-F | GGAAGTACgctTCAATCTACGAGCAATACGCATC |
| PTS-H462K-F | ACGAGAGaaaGTTCGCACTGCAGTAGAATGCTA |
| PTS-Y4L-F | GGAGTTGttgGCCCAAAGTGTTGGAGTGGG |
| PTS-H162K-F | CAGAGTCaaaGGAGAAGACGTCCTTGATAATGCT |
| PTS-A183W-F | tggACTTTGAACGATCCAACCGCG |
| PTS-D187L-F | TTTGAACttgCCAACCGCGAAACAAGTCC |
| PTS-S483K-F | GGCCGTGaaaGAATTCTACAACCAAATGGAGTCAGC |
| PTS-R267L-F | CGTTttgGATCGATTGGTGGAGTCCTACTTC |
| PTS-S278A-F | TGGGCTgctGGATCTTATTTCGAACCGAATTATT |
| PTS-H194L-F | CGCGAAACAAGTCttgAACGCATTGAATGAGTTCTCTTTTC |
| PTS-Q192P-F | GCGAAAccaGTCCACAACGCATTGAATGAGTT |
| PTS-V341E-F | AATAgaaTACAAGGCCCTTTTGGATGTG |
| PTS-E130P-F | ccaAAGTTTAAGGATGGCAAAGATGGA |
| PTS-D250I-F | GGGAGCTGAGTGAAattTCTAGGTGGTGGAAGACTTTACAAGT |
| PTS-D250L-F | GTGAAttgTCTAGGTGGTGGAAGACTTTACAAGT |
| PTS-D250V-F | GTGAAgttTCTAGGTGGTGGAAGACTTTACAAGT |
| PTS-D250M-F | GTGAAatgTCTAGGTGGTGGAAGACTTTACAAGT |
| PTS-D250T-F | GTGAAactTCTAGGTGGTGGAAGACTTTACAAGT |
| PTS-D250A-F | GAGTGAAgctTCTAGGTGGTGGAAGACTTTACAAGT |
| PTS-R464A-F | AGCACGTTgctACTGCAGTAGAATGCTACATGGAAG |
| PTS-D250F-F | GGGAGCTGAGTGAAtttTCTAGGTGGTGGAAGACTTTACAAGT |
| PTS-R464P-F | AGCACGTTccaACTGCAGTAGAATGCTACATGGAAG |
| PTS-Q258D-F | GACTTTAgatGTGCCCACAAAGCTATCATTCG |
| PTS-D250S-F | GTGAAtctTCTAGGTGGTGGAAGACTTTACAAGT |
| PTS-T189L-F | CGATCCAttgGCGAAACAAGTCCACAACGC |
| PTS-D250C-F | GTGAAtgtTCTAGGTGGTGGAAGACTTTACAAGT |
| PTS-Q544S-F | GtctTTGTACCTTCACCCTGTTCCATATT |
| PTS-D137G-F | GATGGCAAAggtGGATTTAAGGTTCCAAATGAGGAT |
| PTS-A390Y-F | GGAGGGAGCAAAAGtatAAACCCACAACCAAGGAGTATATGA |
| PTS-Y221D-F | CGAGCAAgatGCATCTCATCACAAAGGCTTGC |
| PTS-R464S-F | GCACGTTtctACTGCAGTAGAATGCTACATGGAAG |
| PTS-S280V-F | GGCTTCGGGAgttTATTTCGAACCGAATTATTCGGT |
| PTS-R464V-F | GCACGTTgttACTGCAGTAGAATGCTACATGGAAG |
| PTS-Y221E-F | CGAGCAAgaaGCATCTCATCACAAAGGCTTGC |
| PTS-D250Y-F | GGGAGCTGAGTGAAtatTCTAGGTGGTGGAAGACTTTACAAGT |
| PTS-A361K-F | GCTAGGCaaaCCATATCGAGCCTACTATGGAAAA |
| PTS-G477S-F | GCACAAAGTGtctAAGCAAGAGGCCGTGTCTGA |
| PTS-N169E-F | CGTCCTTGATgaaGCTTTTGACTTCACTAGGAACTACTTG |
| PTS-G227T-F | actTTGCTCAAACTTGCTAAGCTGG |
| PTS-M107L-F | TCATGACttgTACGCCACTGCTCTCAGCTTT |
| PTS-V83E-F | TgaaGAAGATGTTGATGAAGCTTTGAAGA |
| PTS-C23S-F | TCATCCAtctGTGTGGGGAGACAAATTCATTGT |
| PTS-Y552L-F | CCAttgTAAATCCGCTCTAACCGAAAAGG |
| PTS-G368A-F | CGAGCCTACTATgctAAAGAAGCCATGAAATACGCCG |
| PTS-F437I-F | ACCTCCTattATCGAGGCTACATTAATCATTGCC |
| PTS-R464T-F | GCACGTTactACTGCAGTAGAATGCTACATGGAAG |
| PTS-D250Q-F | GTGAAcaaTCTAGGTGGTGGAAGACTTTACAAGT |
| PTS-A361R-F | GCTAGGCagaCCATATCGAGCCTACTATGGAAAA |
| PTS-Y63P-F | TCAGACAACccaATGCGGCAACTGAAAATGGT |
| PTS-F506V-F | ACAGAAgttCCAATCCCTCTACTTTATCTTATTCTCA |
| PTS-Y546L-F | GCAGTTGttgCTTCACCCTGTTCCATATTAAATCC |
| PTS-F153L-F | GAAttgTTCGAAGCCACGCATCTCA |
| PTS-D250N-F | GGGAGCTGAGTGAAaatTCTAGGTGGTGGAAGACTTTACAAGT |
| PTS-H225N-F | CATCTCATaatAAAGGCTTGCTCAAACTTGCTAA |
| PTS-T256D-F | GTGGTGGAAGgatTTACAAGTGCCCACAAAGCTATCA |
| PTS-Q539K-F | GGGTCCTGCAATGaaaAACATCATCAAGCAGTTGTACCTTCA |
| PTS-R464G-F | AGCACGTTggtACTGCAGTAGAATGCTACATGGAAG |
| PTS-G477T-F | GCACAAAGTGactAAGCAAGAGGCCGTGTCTGA |
| PTS-G146D-F | TGAGGATgatGCGGTTGCAGTCCTTGAATT |
| PTS-N285Q-F | CGAACCGcaaTATTCGGTAGCTAGGATGATTTTAGC |
| PTS-E55M-F | AGAatgCTGAAGGAAGCATCAGACAACTACA |
| PTS-A390H-F | AGcatAAACCCACAACCAAGGAGTATATGA |
| PTS-M318L-F | CAAttgTTCACAGATGCAATCGAAAGGTG |
| PTS-E381C-F | TTACATGtgtGAGGCCCAATGGAGGGAGC |
| PTS-E426C-F | GACCAAAtgtGCCTTCGATTGGGTGTTCTCC |
| PTS-D495Y-F | GTCAGCATGGAAGtatATTAATGAGGGGTTCCTCAGACC |
| PTS-N104V-F | gttCATGACATGTACGCCACTGCTC |
| PTS-D98K-F | TGTTTaaaGCTTTCTGCAAGAATAATCATGACA |
| PTS-R377K-F | GCCGCGaaaGCTTACATGGAAGAGGCCCA |
| PTS-L510P-F | CCAATCCCTccaCTTTATCTTATTCTCAATTCAGTCCGA |
| PTS-D27C-F | GTGTGTGGGGAtgtAAATTCATTGTCTACAACCCACAATC |
| PTS-F139E-F | GATGGAgaaAAGGTTCCAAATGAGGATGGAGC |
| PTS-A401C-F | GAAGCTGtgtACCAAGACATGTGGCTACATAACTCT |
| PTS-N449L-F | GGCTCGTCttgGATATTACAGGACACGAGTTTGAGAAA |
| PTS-E44D-F | GAGAGAAgatGCTGAGGAGCTGAAAGTGGAGC |
| PTS-A309E-F | GTGTATGATgaaTATGGTACTTTTGAGGAATTACAAATGT |
| PTS-C126W-F | CAGAGTTTCAtggGAAGTTTTTGAAAAGTTTAAGGATGG |
| PTS-A383W-F | GGAAGAGtggCAATGGAGGGAGCAAAAGGC |
| PTS-R42Y-F | TGGAGAGtatGAAGAGGCTGAGGAGCTGAAAG |
| PTS-H474T-F | GGAAGAGactAAAGTGGGGAAGCAAGAGGC |
| PTS-E473D-F | GGAAgatCACAAAGTGGGGAAGCAAGAGG |
| PTS-P260Y-F | ACAAGTGtatACAAAGCTATCATTCGTTAGAGATCG |
| PTS-I30Q-F | TTCcaaGTCTACAACCCACAATCATGCCA |
| PTS-E272A-F | ATTGGTGgctTCCTACTTCTGGGCTTCGGG |
| PTS-L206Y-F | AAGAGGAtatCCACGCGTGGAAGCAAGG |
| PTS-T256V-F | GTGGTGGAAGgttTTACAAGTGCCCACAAAGCTATCA |
| PTS-Q119P-F | TCTCAGAccaCATGGATACAGAGTTTCATGTGAAGTT |
| PTS-L359Q-F | TGATCAAGcaaGGCGCACCATATCGAGCC |
| PTS-A375D-F | GAAATACgatGCGAGAGCTTACATGGAAGAGG |
| PTS-E155S-F | ATTCTTCtctGCCACGCATCTCAGAGTCCA |
| PTS-L316P-F | TGAGGAAccaCAAATGTTCACAGATGCAATCGA |
| PTS-I408Q-F | GTGGCTACcaaACTCTAATAATATTATCATGTCTTGGAGTGG |
| PTS-T465A-F | TTCGCgctGCAGTAGAATGCTACATGGAAGAGC |
| PTS-I73L-F | GGATGCAttgCAACGATTAGGCATTGACTATCTTTT |
| PTS-L116Q-F | CTTTCGCcaaCTCAGACAACATGGATACAGAGTTTCA |
| PTS-E283I-F | CattCCGAATTATTCGGTAGCTAGGATG |
| PTS-S530Y-F | GGGCGATtatTATACACACGTGGGTCCTGCAA |
| PTS-R204W-F | TTTTCGAtggGGATTGCCACGCGTGGAA |
| PTS-R245P-F | TTGCACAGAccaGAGCTGAGTGAAGATTCTAGGTGGT |
| PTS-T424K-F | GGGCATTGTGaaaAAAGAAGCCTTCGATTGGGTG |
| PTS-F282C-F | GATCTTATtgtGAACCGAATTATTCGGTAGCTAGG |
| PTS-K102P-F | GATGCTTTCTGCccaAATAATCATGACATGTACGCCACTG |
| PTS-A45K-F | AGAAGAGaaaGAGGAGCTGAAAGTGGAGCTGA |
| PTS-K191R-F | ATCCAACCGCGagaCAAGTCCACAACGCATTGAATG |
| PTS-R175K-F | CACTaaaAACTACTTGGAATCAGTCTATGCAACTT |
| PTS-E249P-F | GCTGAGTccaGATTCTAGGTGGTGGAAGACTTTACA |
| PTS-N497Q-F | GGACATTcaaGAGGGGTTCCTCAGACCAACA |
| PTS-A13R-F | TGGGTGCTagaTCTCGTCCTCTTGCGAATTTTC |
| PTS-L331Y-F | GGATGCTTCATGTtatGATAAACTTCCAGATTACATGAAAATAGTATAC |
| PTS-E473A-F | CATGGAAgctCACAAAGTGGGGAAGCAAGAGG |
| PTS-D85H-F | TGTGGAAcatGTTGATGAAGCTTTGAAGAATCTGTT |
| PTS-R118H-F | CCTTCTCcatCAACATGGATACAGAGTTTCATGTGA |
| PTS-D353F-F | GGAAGTTtttGAGGAGTTGATCAAGCTAGGCG |
| PTS-V193Y-R | GTGataTTGTTTCGCGGTTGGATCG |
| PTS-V193S-R | TGTGagaTTGTTTCGCGGTTGGATCG |
| PTS-H462F-R | TGCGAACaaaCTCTCGTTTTTTCTCAAACTCGTG |
| PTS-V193W-R | CGTTGTGccaTTGTTTCGCGGTTGGATCG |
| PTS-V193R-R | TGTGtctTTGTTTCGCGGTTGGATCG |
| PTS-D187P-R | CGGTTGGtggGTTCAAAGTTGCATAGACTGATTCCA |
| PTS-G499F-R | GGTCTGAGGAAaaaCTCATTAATGTCCTTCCATGCTGA |
| PTS-V288M-R | CATCCTAGCcatCGAATAATTCGGTTCGAAATAAGA |
| PTS-Q488L-R | CTGACTCCATcaaGTTGTAGAATTCAGACACGGCCT |
| PTS-L178P-R | CTGATTCtggGTAGTTCCTAGTGAAGTCAAAAGCATTA |
| PTS-A183H-R | GGATCGTTCAAAGTatgATAGACTGATTCCAAGTAGTTCCTAGTGA |
| PTS-L56Y-R | GCTTCCTTataCTCTCTTTTCAGCTCCACTTTCAGC |
| PTS-A241F-R | CCCTTCTGTGCAAaaaTTGTACCAAGTTGAAATCCAGCTT |
| PTS-L56W-R | CTTCCTTccaCTCTCTTTTCAGCTCCACTTTCAGC |
| PTS-H194K-R | tttGACTTGTTTCGCGGTTGGATC |
| PTS-E46P-R | TTTCAGCTCtggAGCCTCTTCTCTCTCTCCAGCC |
| PTS-A241I-R | CCCTTCTGTGCAAaatTTGTACCAAGTTGAAATCCAGCTT |
| PTS-H225Y-R | GCCTTTataATGAGATGCGTATTGCTCGTAGAT |
| PTS-G146L-R | CAACCGCcaaATCCTCATTTGGAACCTTAAATCC |
| PTS-P362F-R | CTCGATAaaaTGCGCCTAGCTTGATCAACTC |
| PTS-V83W-R | CATCAACATCTTCccaAAAAAGATAGTCAATGCCTAATCGTT |
| PTS-Q488M-R | CTGACTCCATcatGTTGTAGAATTCAGACACGGCCT |
| PTS-Y367L-R | CTTTTCCcaaGTAGGCTCGATATGGTGCGC |
| PTS-H225K-R | GCCTTTtttATGAGATGCGTATTGCTCGTAGAT |
| PTS-V467L-R | AGCATTCcaaTGCAGTGCGAACGTGCTCT |
| PTS-G499C-R | GTCTGAGGAAacaCTCATTAATGTCCTTCCATGCTGA |
| PTS-A309P-R | CCATAtggATCATACACATCATCCATAAGAGATAATACA |
| PTS-K478W-R | TCTTGccaCCCCACTTTGTGCTCTTCCA |
| PTS-P188M-R | TCGCGGTcatATCGTTCAAAGTTGCATAGACTGATT |
| PTS-M372S-R | GCGTATTTagaGGCTTCTTTTCCATAGTAGGCTCG |
| PTS-R267P-R | TCGATCtggAACGAATGATAGCTTTGTGGGC |
| PTS-V193Q-R | TGTGttgTTGTTTCGCGGTTGGATCG |
| PTS-V10F-R | CAGCACCaaaTCCAACACTTTGGGCATACAAC |
| PTS-S14V-R | AAGAGGACGaacAGCAGCACCCACTCCAACAC |
| PTS-A156Y-R | GATGCGTataTTCGAAGAATTCAAGGACTGCA |
| PTS-P335W-R | CATGTAATCccaAAGTTTATCTAAACATGAAGCATCCC |
| PTS-R204L-R | GGCAATCCcaaTCGAAAAGAGAACTCATTCAATGC |
| PTS-I215A-R | GATTGAagcGTACTTCCTTGCTTCCACGCG |
| PTS-H462K-R | TGCGAACtttCTCTCGTTTTTTCTCAAACTCGTG |
| PTS-Y4L-R | TTTGGGCcaaCAACTCCATGTTTTTTCTCCTTGA |
| PTS-H162K-R | CTTCTCCtttGACTCTGAGATGCGTGGCTTCG |
| PTS-A183W-R | GGATCGTTCAAAGTccaATAGACTGATTCCAAGTAGTTCCTAGTGA |
| PTS-D187L-R | CGGTTGGcaaGTTCAAAGTTGCATAGACTGATTCCA |
| PTS-S483K-R | AGAATTCtttCACGGCCTCTTGCTTCCC |
| PTS-R267L-R | CCAATCGATCcaaAACGAATGATAGCTTTGTGGGC |
| PTS-S278A-R | TAAGATCCagcAGCCCAGAAGTAGGACTCCACC |
| PTS-H194L-R | TcaaGACTTGTTTCGCGGTTGGATC |
| PTS-Q192P-R | TTGTGGACtggTTTCGCGGTTGGATCGTTC |
| PTS-V341E-R | GGGCCTTGTAttcTATTTTCATGTAATCTGGAAGTTTATCTAAA |
| PTS-E130P-R | CCATCCTTAAACTTtggAAAAACTTCACATGAAACTCTGTATCC |
| PTS-D250I-R | aatTTCACTCAGCTCCCTTCTGTGC |
| PTS-D250L-R | CCACCTAGAcaaTTCACTCAGCTCCCTTCTGTGC |
| PTS-D250V-R | CCACCTAGAaacTTCACTCAGCTCCCTTCTGTGC |
| PTS-D250M-R | CCACCTAGAcatTTCACTCAGCTCCCTTCTGTGC |
| PTS-D250T-R | CCACCTAGAagtTTCACTCAGCTCCCTTCTGTGC |
| PTS-D250A-R | ACCTAGAagcTTCACTCAGCTCCCTTCTGTGC |
| PTS-R464A-R | TGCAGTagcAACGTGCTCTCGTTTTTTCTCAA |
| PTS-D250F-R | aaaTTCACTCAGCTCCCTTCTGTGC |
| PTS-R464P-R | TGCAGTtggAACGTGCTCTCGTTTTTTCTCAA |
| PTS-Q258D-R | TGGGCACatcTAAAGTCTTCCACCACCTAGAATCTTC |
| PTS-D250S-R | CCACCTAGAagaTTCACTCAGCTCCCTTCTGTGC |
| PTS-T189L-R | GTTTCGCcaaTGGATCGTTCAAAGTTGCATAGA |
| PTS-D250C-R | CCACCTAGAacaTTCACTCAGCTCCCTTCTGTGC |
| PTS-Q544S-R | GGTGAAGGTACAAagaCTTGATGATGTTTTGCATTGCAG |
| PTS-D137G-R | AATCCaccTTTGCCATCCTTAAACTTTTCAAA |
| PTS-A390Y-R | ataCTTTTGCTCCCTCCATTGGG |
| PTS-Y221D-R | GAGATGCatcTTGCTCGTAGATTGATATGTACTTCCT |
| PTS-R464S-R | CTGCAGTagaAACGTGCTCTCGTTTTTTCTCAA |
| PTS-S280V-R | AATAaacTCCCGAAGCCCAGAAGTAGG |
| PTS-R464V-R | CTGCAGTaacAACGTGCTCTCGTTTTTTCTCAA |
| PTS-Y221E-R | GAGATGCttcTTGCTCGTAGATTGATATGTACTTCCT |
| PTS-D250Y-R | ataTTCACTCAGCTCCCTTCTGTGC |
| PTS-A361K-R | GATATGGtttGCCTAGCTTGATCAACTCCTCG |
| PTS-G477S-R | GCTTagaCACTTTGTGCTCTTCCATGTAGCA |
| PTS-N169E-R | AAGCttcATCAAGGACGTCTTCTCCATGG |
| PTS-G227T-R | GCAAGTTTGAGCAAagtTTTGTGATGAGATGCGTATTGCT |
| PTS-M107L-R | TGGCGTAcaaGTCATGATTATTCTTGCAGAAAGCA |
| PTS-V83E-R | CATCAACATCTTCttcAAAAAGATAGTCAATGCCTAATCGTT |
| PTS-C23S-R | CCCACACagaTGGATGAAAATTCGCAAGAGG |
| PTS-Y552L-R | GAGCGGATTTAcaaTGGAACAGGGTGAAGGTACAACT |
| PTS-G368A-R | TTagcATAGTAGGCTCGATATGGTGCGC |
| PTS-F437I-R | CCTCGATaatAGGAGGTCGGGAGAACACCC |
| PTS-R464T-R | CTGCAGTagtAACGTGCTCTCGTTTTTTCTCAA |
| PTS-D250Q-R | CCACCTAGAttgTTCACTCAGCTCCCTTCTGTGC |
| PTS-A361R-R | GATATGGtctGCCTAGCTTGATCAACTCCTCG |
| PTS-Y63P-R | CGCATtggGTTGTCTGATGCTTCCTTCAGCT |
| PTS-F506V-R | GGGATTGGaacTTCTGTTGGTCTGAGGAACCCC |
| PTS-Y546L-R | GGTGAAGcaaCAACTGCTTGATGATGTTTTGCA |
| PTS-F153L-R | GTGGCTTCGAAcaaTTCAAGGACTGCAACCGCTC |
| PTS-D250N-R | attTTCACTCAGCTCCCTTCTGTGC |
| PTS-H225N-R | GCCTTTattATGAGATGCGTATTGCTCGTAGAT |
| PTS-T256D-R | GTAAatcCTTCCACCACCTAGAATCTTCACTC |
| PTS-Q539K-R | TtttCATTGCAGGACCCACGTGTG |
| PTS-R464G-R | TGCAGTaccAACGTGCTCTCGTTTTTTCTCAA |
| PTS-G477T-R | GCTTagtCACTTTGTGCTCTTCCATGTAGCA |
| PTS-G146D-R | CAACCGCatcATCCTCATTTGGAACCTTAAATCC |
| PTS-N285Q-R | CCGAATAttgCGGTTCGAAATAAGATCCCGA |
| PTS-E55M-R | GCTTCCTTCAGcatTCTTTTCAGCTCCACTTTCAGCT |
| PTS-A390H-R | GGTTGTGGGTTTatgCTTTTGCTCCCTCCATTGGG |
| PTS-M318L-R | GCATCTGTGAAcaaTTGTAATTCCTCAAAAGTACCATATGC |
| PTS-E381C-R | GGGCCTCacaCATGTAAGCTCTCGCGGCG |
| PTS-E426C-R | CGAAGGCacaTTTGGTCACAATGCCCTCTTCC |
| PTS-D495Y-R | TataCTTCCATGCTGACTCCATTTGG |
| PTS-N104V-R | GCGTACATGTCATGaacATTCTTGCAGAAAGCATCAAACAT |
| PTS-D98K-R | GCAGAAAGCtttAAACATTTCAAACAGATTCTTCAAAGC |
| PTS-R377K-R | ATGTAAGCtttCGCGGCGTATTTCATGGC |
| PTS-L510P-R | TAAAGtggAGGGATTGGAAATTCTGTTGGTC |
| PTS-D27C-R | TTTacaTCCCCACACACATGGATGAAA |
| PTS-F139E-R | GGAACCTTttcTCCATCTTTGCCATCCTTAAACT |
| PTS-A401C-R | TCTTGGTacaCAGCTTCATATACTCCTTGGTTGTG |
| PTS-N449L-R | AATATCcaaGACGAGCCTGGCAATGATTAA |
| PTS-E44D-R | CCTCAGCatcTTCTCTCTCTCCAGCCTGGCA |
| PTS-A309E-R | CCATAttcATCATACACATCATCCATAAGAGATAATACA |
| PTS-C126W-R | CTTCccaTGAAACTCTGTATCCATGTTGTCTGA |
| PTS-A383W-R | TCCATTGccaCTCTTCCATGTAAGCTCTCGCG |
| PTS-R42Y-R | CCTCTTCataCTCTCCAGCCTGGCATGATTG |
| PTS-H474T-R | CCACTTTagtCTCTTCCATGTAGCATTCTACTGCA |
| PTS-E473D-R | CCACTTTGTGatcTTCCATGTAGCATTCTACTGCAGTG |
| PTS-P260Y-R | GCTTTGTataCACTTGTAAAGTCTTCCACCACCTAG |
| PTS-I30Q-R | GGGTTGTAGACttgGAATTTGTCTCCCCACACACATG |
| PTS-E272A-R | AGTAGGAagcCACCAATCGATCTCTAACGAATGA |
| PTS-L206Y-R | CGCGTGGataTCCTCTTCGAAAAGAGAACTCATTC |
| PTS-T256V-R | GTAAaacCTTCCACCACCTAGAATCTTCACTC |
| PTS-Q119P-R | ATCCATGtggTCTGAGAAGGCGAAAGCTGAGA |
| PTS-L359Q-R | TGCGCCttgCTTGATCAACTCCTCGTCAACTTC |
| PTS-A375D-R | CTCTCGCatcGTATTTCATGGCTTCTTTTCCATAGT |
| PTS-E155S-R | GCGTGGCagaGAAGAATTCAAGGACTGCAACCG |
| PTS-L316P-R | ACATTTGtggTTCCTCAAAAGTACCATATGCATCA |
| PTS-I408Q-R | TAGAGTttgGTAGCCACATGTCTTGGTTGCC |
| PTS-T465A-R | TTCTACTGCagcGCGAACGTGCTCTCGTTTTT |
| PTS-I73L-R | ATCGTTGcaaTGCATCCACCATTTTCAGTTGC |
| PTS-L116Q-R | GTCTGAGttgGCGAAAGCTGAGAGCAGTGG |
| PTS-E283I-R | CCGAATAATTCGGaatGAAATAAGATCCCGAAGCCCA |
| PTS-S530Y-R | GTGTATAataATCGCCCTCTTTGTAAATAACCTC |
| PTS-R204W-R | GCAATCCccaTCGAAAAGAGAACTCATTCAATGC |
| PTS-R245P-R | AGCTCtggTCTGTGCAAAGCTTGTACCAAGTT |
| PTS-T424K-R | CTTTtttCACAATGCCCTCTTCCACTCC |
| PTS-F282C-R | CGGTTCacaATAAGATCCCGAAGCCCAGAA |
| PTS-K102P-R | TTtggGCAGAAAGCATCAAACATTTCAA |
| PTS-A45K-R | GCTCCTCtttCTCTTCTCTCTCTCCAGCCTGGC |
| PTS-K191R-R | TTGtctCGCGGTTGGATCGTTCAA |
| PTS-R175K-R | CCAAGTAGTTtttAGTGAAGTCAAAAGCATTATCAAGGA |
| PTS-E249P-R | TAGAATCtggACTCAGCTCCCTTCTGTGCAAA |
| PTS-N497Q-R | ACCCCTCttgAATGTCCTTCCATGCTGACTCC |
| PTS-A13R-R | ACGAGAtctAGCACCCACTCCAACACTTTGG |
| PTS-L331Y-R | CataACATGAAGCATCCCACCTTTCG |
| PTS-E473A-R | CTTTGTGagcTTCCATGTAGCATTCTACTGCAGTG |
| PTS-D85H-R | CATCAACatgTTCCACAAAAAGATAGTCAATGCC |
| PTS-R118H-R | CATGTTGatgGAGAAGGCGAAAGCTGAGAGC |
| PTS-D353F-R | ACTCCTCaaaAACTTCCTCAAACACATCCAAAAGG |
| 1114a up-F | GAGAAATGTTGGGATCCAG |
| 1114a up-pGAL1-R | ATTACTGAAAGTTCCAAAGAGAAGGGATAATAGTACAAACTTACATAGCGT |
| 1114a down-tCYC1-F | TGAGAAGGTTTTGGGACGCTCGAAGAGGACAGTCAATAGCATCAT |
| 1114a down-R | AGATAAGAAGTGGGAAGGTAAAATC |
| 1114a val-F | TGTATTTGATCTTCCATGTGC |
| 1114a val-R | TCATAGTATCGATGACTCAGATG |
